# CUSTOM-Muscle: Customisable, Shape-Tuneable, Oriented Modular PDMS Scaffolds for Disease-Relevant 3D Skeletal Muscle Engineering

**DOI:** 10.64898/2026.11.12.766110

**Authors:** Eugenia Carraro, Daniel Moore, Sungwoo Choi, Federico Nebuloni, Taylor Minckley, Sudeep Joshi, Albane Imbert, Francesco Saverio Tedesco, Andrea Serio

## Abstract

Modelling skeletal muscle *in vitro* offers a powerful approach for studying muscle physiology and disease. Despite their potential, current models offer limited scope to combine customisable three-dimensional architecture and mechanical support with complementary functional readouts. Many also lack key features of native muscle, including cellular maturation, neuromuscular organisation and a physiological mechanical environment, limiting their ability to recapitulate muscle structure and function. Here, we establish a three-dimensional (3D) miniaturised, customisable and multifunctional platform for human neuromuscular tissue engineering. We developed an entirely open-source pipeline combining computer-aided design (CAD), stereolithography (SLA) and fused deposition modelling (FDM) to generate culture platforms with tuneable 3D geometries and mechanical properties, supporting short- and long-term culture. Across the configurations tested, all platforms supported skeletal muscle differentiation and functional assessment of twitch and tetanic forces. Integration with motor neuron cultures enabled the generation of neuromuscular constructs suitable for live imaging, metabolic analyses and identification of putative neuromuscular contact sites. Furthermore, the platform recapitulated aberrant nuclear elongation associated with muscle laminopathies, enabling quantitative assessment of disease-associated nuclear morphology. Finally, to address the critical role of mechanical stimulation in muscle physiology, we designed and fabricated an integrated platform for controlled mechanical elongation of 3D skeletal muscle constructs. By combining architectural and mechanical customisation with complementary functional readouts, this framework enables experiments tailored to how tissue organisation, intercellular interactions and external mechanical cues shape muscle function and disease. Its open-source foundation provides a route to wider adoption and adaptation across tissue engineering and the biosciences.

**Graphical abstract:** 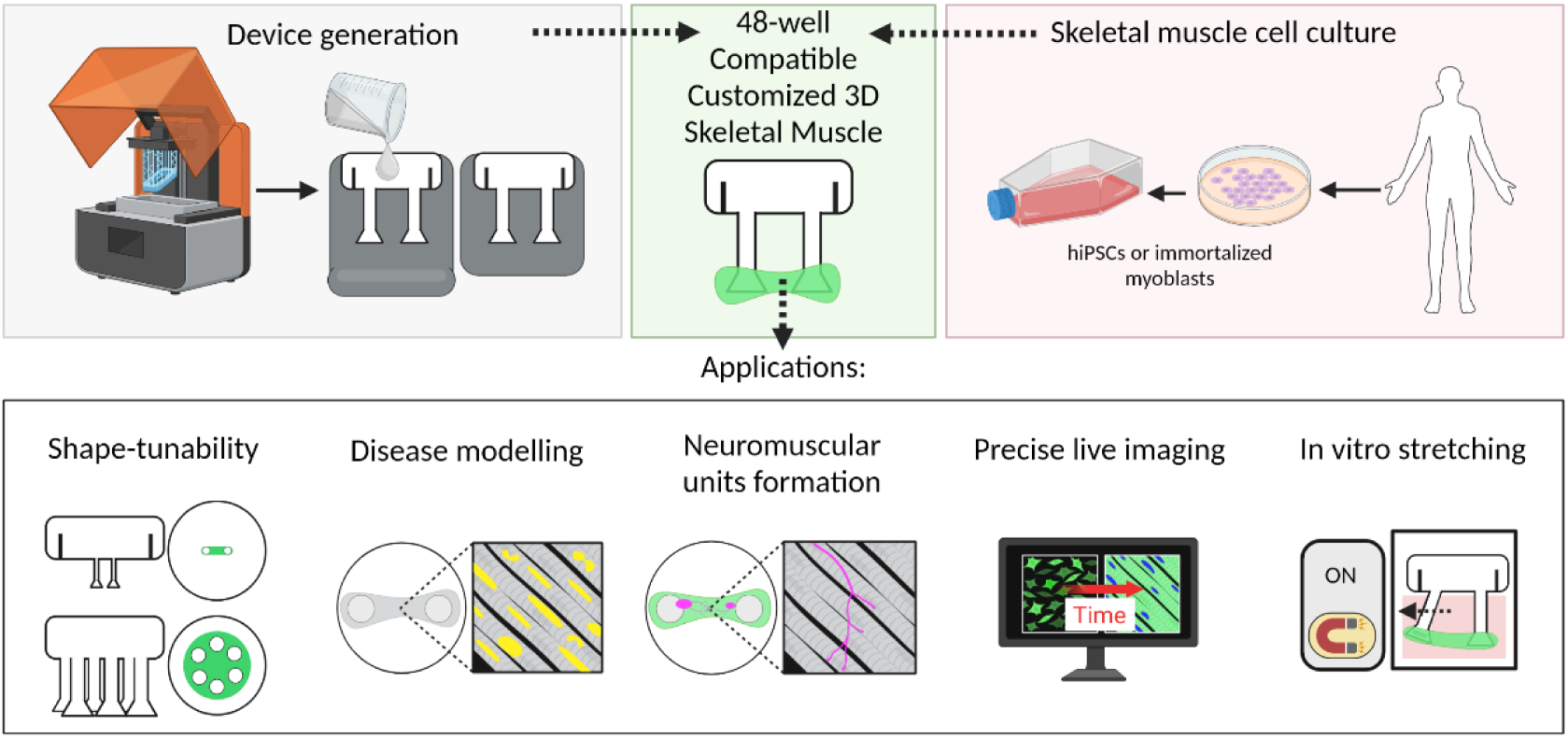

## Introduction

Skeletal muscle is a highly complex tissue composed of multiple cell types and is responsible for essential physiological functions, including locomotion and respiration. The homeostasis of skeletal muscle tissue relies on a finely tuned balance of repetitive regeneration cycles, during which resident muscle stem cells, known as satellite cells, commit into myoblasts and fuse either with pre-existing myofibers or with one another to regenerate damaged tissue^1^. In addition, several other cell types contribute to the overall health and functionality of the tissue, including, motor neurons, which are essential for promoting physiological muscle contraction. Skeletal muscle is the target of a wide range of incurable diseases affecting infant, paediatric, and adult patients. Generally classified as inherited or acquired, these diseases can compromise skeletal muscle integrity, as observed in volumetric muscle loss, disrupt sarcolemma stability, as in muscular dystrophies, or impair nuclear integrity, as in laminopathies^2^. Despite extensive efforts over the past decades to develop and employ multiple animal models, not only to elucidate disease hallmarks but also to evaluate therapeutic strategies, effective treatments remain largely unavailable. While in vivo models have played a critical role in uncovering pathophysiological mechanisms, they are expensive, time-consuming, and unsuitable for high-throughput early-stage studies. By comparison, human cell–based *in vitro* models offer a more accessible approach for investigating human-specific biological processes. Furthermore, due to increasingly stringent ethical constraints, the need for alternative experimental systems to replace or reduce animal models has become increasingly evident. In this context, the possibility of recreating skeletal muscle tissue *in vitro* has been extensively investigated in recent years through the application of multiple experimental techniques ranging from microfluidics to 3D printing technologies or to the generation of three-dimensional elongated gels anchored to diverse supporting platforms^3–7^. Overall, generating miniaturised yet sufficiently large functional skeletal muscle tissues represent an important challenge in tissue engineering, requiring the integration of multiple strategies that account for cellular diversity, biomaterials, and external biophysical and biomechanical cues. Maintaining an optimal tissue size is essential to balance experimental manipulation and throughput with sufficient tissue volume and myofiber numbers, while enabling the production of constructs that can be readily scaled up or down for different experimental applications.

Morphological and functional features of humanized skeletal muscle can be recapitulated *in vitro* through various approaches, including micropatterning ^8,9^, organoid generation^10^, or combining decellularization with recellularization ^11,12^. While these strategies are promising, they are limited by challenges such as restricted tissue size, difficulties in manipulation, dependence on donor material, and the lack of precise and controllable mechanical cues. In particular, engineered skeletal muscle constructs must be subjected to appropriate tensile forces and compensate for the absence of native supporting structures such as tendons and bones. Hence, most current approaches, including ours, rely on multistep fabrication of biocompatible polydimethylsiloxane (PDMS) supports incorporating two or more T-shaped pillars. These structures provide mechanical anchoring, enable controlled tension during tissue maturation and promote muscle fibre alignment. However, conventional platforms impose constraints on both tissue scale and customisation. Constructs must be sufficiently large to capture the cellular and structural complexity of skeletal muscle and permit practical handling, yet sufficiently small to limit cell and reagent consumption and support scalable production. More importantly, standardised support geometries offer limited control over tissue shape and mechanical environment, despite the substantial anatomical and biomechanical diversity of skeletal muscles. This limitation is particularly relevant to disease modelling, as different muscles exhibit distinct structural features and disease susceptibilities that may require tissue-specific geometries and mechanical cues^13,14^. In parallel, tissue fabrication, maturation and functional assessment are frequently performed using separate platforms, further limiting experimental flexibility and throughput. We therefore asked whether a customisable, vertically integrated ecosystem of smart devices could overcome these limitations by enabling skeletal muscle constructs with tuneable geometries and mechanical properties, while integrating tissue culture, maturation and functional assessment within a unified experimental framework.

Building on our previous results^9^, we hypothesize that the development of a reliable, miniaturized and user-friendly pipeline that accounts for the complexity of skeletal muscle tissue could provide the appropriate cues for prolonged 3D *in vitro* culture. Such a platform would also serve as a versatile tool for disease modelling, live imaging, and co-culture experiments. Thus, here, we present a combination of stereolithography (SLA) and fused deposition modelling (FDM) printing techniques as a simple, open-source platform for the in-lab fabrication of a wide range of biocompatible devices. These devices can be readily adapted in size and shape and tuned to exhibit distinct mechanical properties, making them suitable for studying multiple aspects of skeletal muscle tissue.

Furthermore, rather than reducing skeletal muscle complexity to facilitate *in vitro* manipulation, we sought to preserve key structural and functional features of native muscle within a controllable and scalable system. To this end, we integrated these bioengineering platforms with human induced pluripotent stem cell (hiPSC)-derived skeletal muscle cells to establish advanced *in vitro* models that better capture the architecture and function complexity of skeletal muscle. Using this approach, we successfully recapitulated characteristic histo-pathological features of *LMNA*-related muscular dystrophy, demonstrating the capacity of the system to model disease-specific alterations under controlled conditions. In addition, this platform offers high modularity and adaptability, allowing force measurement, live imaging and metabolic analysis to be integrated within the same experimental framework and applied across diverse experimental settings. We further demonstrate the compatibility of this platform with motor neuron (MN) co-culture, supporting the generation of neuromuscular constructs within the same engineered muscle microenvironment, providing a tractable system to investigate muscle-neuron interactions. Unlike approaches that use separate chambers or compartmentalised microfluidic systems to spatially organise muscle and motor neurons, our platform enables their direct integration within a single three-dimensional construct, providing a tractable system for investigating muscle–neuron interactions and neuromuscular junction (NMJ) formation. Coupling the platform with high-resolution live imaging further enables continuous, non-invasive monitoring of tissue morphology, fibre alignment and contractile behaviour throughout culture.

Finally, we introduce a versatile stimulation and conditioning system capable of applying programmable and reproducible skeletal muscle elongation and contraction *in vitro*. By enabling controlled mechanical training and dynamic tissue actuation, this platform provides a powerful tool for studying skeletal muscle development, maturation, mechanobiology, and disease progression, while also expanding opportunities for drug screening, regenerative medicine, and biohybrid tissue engineering applications.

## Results

### Combination of resin and ASA printing techniques for the generation and validation of 48-well PDMS skeletal muscle supports with tuneable geometry

Taking advantage of our previously established open source fabrication pipeline^9^, we designed and microfabricated skeletal muscle scaffolds differing in size and geometry (Figure 1A). Each support is generated by assembling two mould components, as previously described^9^, which are held tightly together by an elastic TPU (Thermoplastic Polyurethane)-printed ring. Casting uncured PDMS into the assembled moulds, followed by thermal curing and mould demoulding, results in the production of biocompatible PDMS supports. These supports are characterized by the presence of two or multiple pillars, up to six, equipped with conical bases. The versatility of our fabrication approach enables customization of key design parameters that govern both the architecture and mechanical properties of the engineered skeletal muscle. Specifically, the number and special arrangement of the pillars can be readily adjusted, while geometrical features such as pillar diameter, height, and base morphology can be tuned to modulate tissue anchorage, mechanical resistance, and force transmission. This design flexibility allows the platform to be adapted to diverse experimental applications and facilitates the rational engineering of tissue-specific mechanical environments (Supplementary Table 1). We next evaluated the dimensional limits of our approach (Supplementary Figure 1D–E) and generated miniaturized versions of the devices capable of supporting the formation of 3D skeletal muscle constructs of different sizes (∼8.5 mm as well as ∼4.7 mm-long from the external edge of the pillars). For these configurations, we were able to fabricate all components required for hydrogel casting, including the FDM ASA–printed casting systems (Supplementary Figure 1A-C), enabling the generation of viable and *in vitro*–cultivable 3D skeletal muscle tissues (Figure 1B, E). We generated a wide range of support designs (*L-P, M1,* Supplementary Figure 2A-D), including a miniaturized version of the device supporting the formation of “twinned” muscles on the same platform (*M2,* Figure 1A). This configuration enables parallel *in vitro* maintenance of two miniaturized muscle constructs, thereby opening new opportunities for multiplexed experiments, reduced reagent consumption, and the investigation of cell–cell interactions (Figure 1C). Despite the miniaturized dimensions of these supports, the resulting engineered skeletal muscles differentiated robustly (Supplementary Figure 2E-F). Additionally, we explored the possibility of modulating not only the dimensions but also the shape of the *in vitro* 3D constructs. Motivated by the central role of mechanical tension and the diverse architectures of skeletal muscle across the human body and mammalian systems, we extended this pipeline to generate circular 3D skeletal muscle constructs, reminiscent of the radial organisation of the myofibers in the diaphragm. In these configurations, regions of the gels likely experienced distinct tension profiles, as observed by differences in skeletal muscle cell orientation (Figure 1D). These observations are consistent with previous studies demonstrating the role of multiple attachment points in regulating muscle organization^15^.

**Figure 1.**
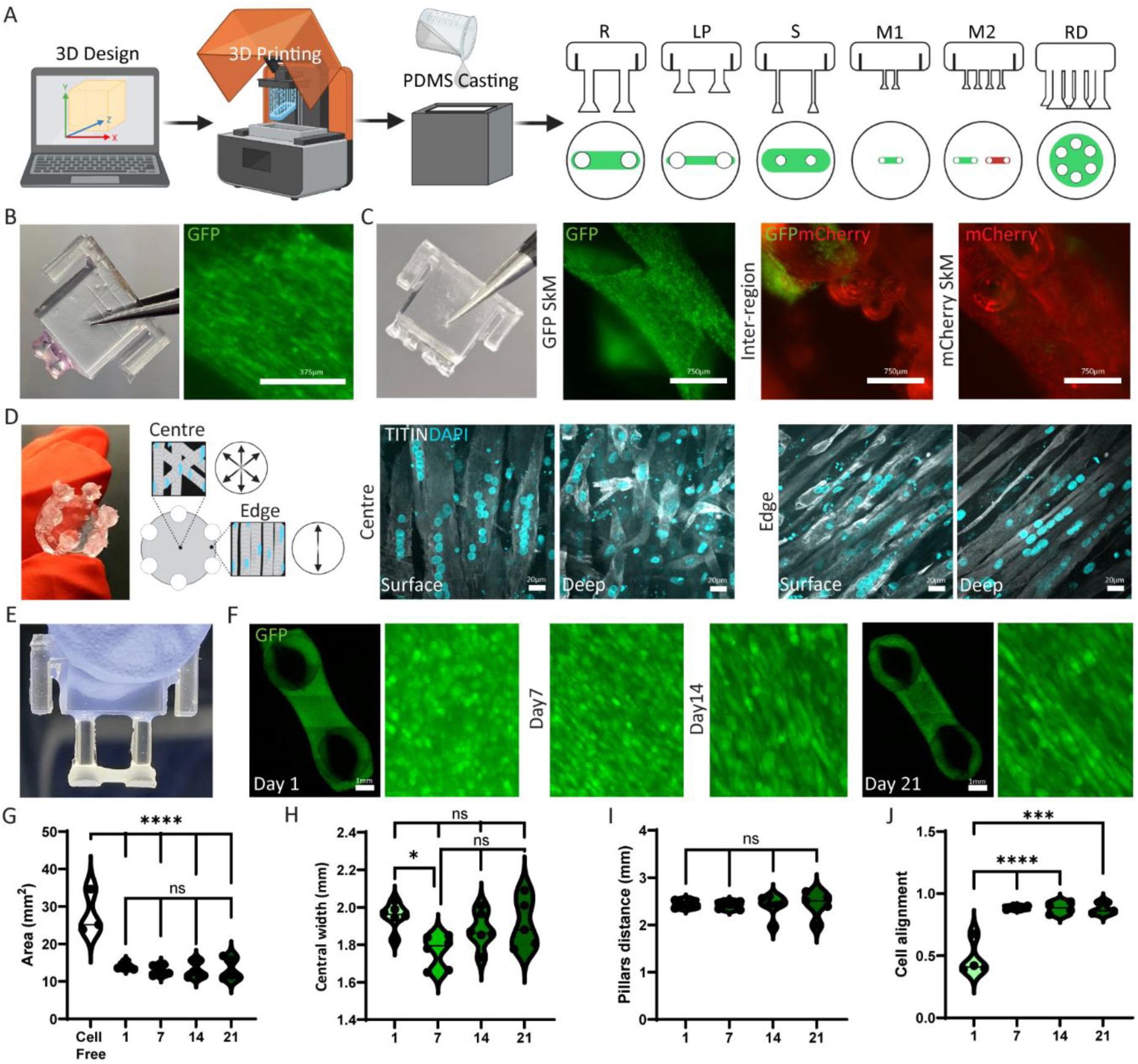
Generation and validation of PDMS scaffolds for miniaturized (48-well) engineered skeletal muscles with tuneable geometry. (A) Graphical abstract of support generation (R reference, L-P low-profile, S slender, M1 miniaturized, M2 miniaturized double, RD radial). (B) M1 muscle representative image and respective SkM tissue differentiation. (C) M2 muscle representative image and respective SkM tissue differentiation with a focus on GFP SkM, inter-region and mCherry SkM. (D) RD representative image and predictive cell orientation and representative SkM tissue differentiation, with a focus on central and peripheral regions. (E) Representative image of 3D SkM on a Reference support. (F) Representative images of a 3D SkM maturation from day 1 to day 21. (G-J) Morphometric analysis of 3D SkM gels at day 1, 7, 14, 21. Scale bars 375, 750, 20µm and 1mm. GFP in green, mCherry in red, TITIN in grey, DAPI in cyan.

We then tested the durability of our platform, assessing long-term maintenance of 3D skeletal muscle constructs. As contractile forces generated by differentiated myofibers can lead to early detachment of cell layers, we monitored constructs generated from immortalized human skeletal muscle cells at multiple time points, up to 28 days in culture (Supplementary Figure 2G, H). Although these cells are typically maintained for approximately 14 days^6,5,16^, extended culture was enabled by the optimized support design. Based on quantitative assessments of constructs area, preservation of central width, and stable inter-pillar distance, together with a healthy, non-punctate appearance of skeletal muscle cells, 21 days was identified as the optimal time point for subsequent analyses, in line with other literature data for late differentiation analysis^17^ (Figure 1G-I). Skeletal muscle architecture is characterized by bundles of parallel-oriented myofibers embedded within a highly organized extracellular matrix (ECM) composed primarily of laminins and collagens^18,19^. In our system, this ECM is substituted with Matrigel and maintained through appropriate media supplementation (Supplementary Figure 2I, J). Particularly in longitudinal muscles, recapitulating this organization represents a key requirement for relevant *in vitro* models. We therefore quantified cell alignment over time (Figure 1J), defined as the degree to which skeletal muscle cells adopted a common orientation corresponding to the axis imposed by the supporting pillars.

Cell alignment increased progressively during culture, until day 7 of differentiation, which coincided with the onset of myoblast differentiation following the switch to differentiation medium, indicating a transition toward a more mature phenotype (Figure 1F).

### Support geometries differentially modulate contractile function in customized muscle constructs while preserving skeletal muscle maturation

Skeletal muscle function is defined by its ability to contract. Detection and quantification of contractile behaviour *in vitro* provided a direct readout of construct viability and functional performance and enabled the comparison of contractile properties across different cell lines, in healthy and pathological conditions. It is well established in the literature that different techniques to generate SkM *in vitro*, can impact the ability to detect contractile activity^20,21^. In particular, an appropriate balance between flexibility and stiffness is required to enable maturation and measurable contraction, thereby constraining both the choice of fabrication materials and the dimensions of the constructs. In this context, we established a pipeline in which, following myoblast expansion and hydrogel generation, *in vitro* contractile activity can be assessed after 21 days in culture (Figure 2A). Using this customized system (Supplementary Figure 3A-C), we compared two PDMS support designs, referred to as *Reference* and *Slender*. While the overall pillar geometry was preserved, the *Slender* supports exhibited a significant reduction in its diameter and corresponding conical base dimensions (Figure 2B, C). This geometric reduction resulted in altered mechanical properties of the supports (Supplementary Table 1). 3D muscles generated with the two PDMS designs were subsequently evaluated for their impact on functionally associated phenotypes, including contractile behaviour. Skeletal muscle constructs displayed spontaneous contractile activity in culture between days 14 and 21. At 21 days post differentiation we assessed two contractile parameters: twitch force, corresponding to rapid contraction-relaxation events, and tetanic force, representing maximal sustained contraction. The experimental setup compared both *Reference* and *Slender* designs and two initial cell densities 1×10⁶ cells per 100µL hydrogel, commonly reported in similar studies^5,22^ and a reduced density of 5×10⁵ cells per 100µL hydrogel, representing distinct microenvironmental conditions. At the higher cell density (1×10⁶ cells), constructs cultured on *Slender* supports exhibited significantly higher tetanic force and increased twitch peak compared to those grown on reference supports. In contrast, at the lower cell density (5×10⁵ cells), no significant differences in contractile parameters were detected between the two support designs. When analysing the effect of cell density within each support type, significant differences between the two cell concentrations were observed only in constructs cultured on *Slender* supports. No comparable differences were detected on *Reference* supports (Figure 2D, E). Collectively, these results indicate that thinner, softer supports enhance the detection of contractile phenotypes and reveal sensitivity to cell density–dependent effects, and, by extension, have the potential to reveal cell line specific contractile properties that are not detectable using stiffer support designs. Interestingly, despite the functional differences observed, robust differentiation was detected across all four experimental conditions evaluated (Figure 2F-I). Support design did not affect myogenic differentiation, as assessed by the expression of the late myogenic marker TITIN; differences in its coverage were observed only as a consequence of reduced cell density (Figure 2J). In parallel, no significant differences were detected in myotube formation (Figure 2K), nor in sarcomere organization (Figure 2M). Conversely, myotube thickness was influenced by the scaffold properties and cell density, suggesting differences among the four conditions tested (Figure 2L).

**Figure 2.**
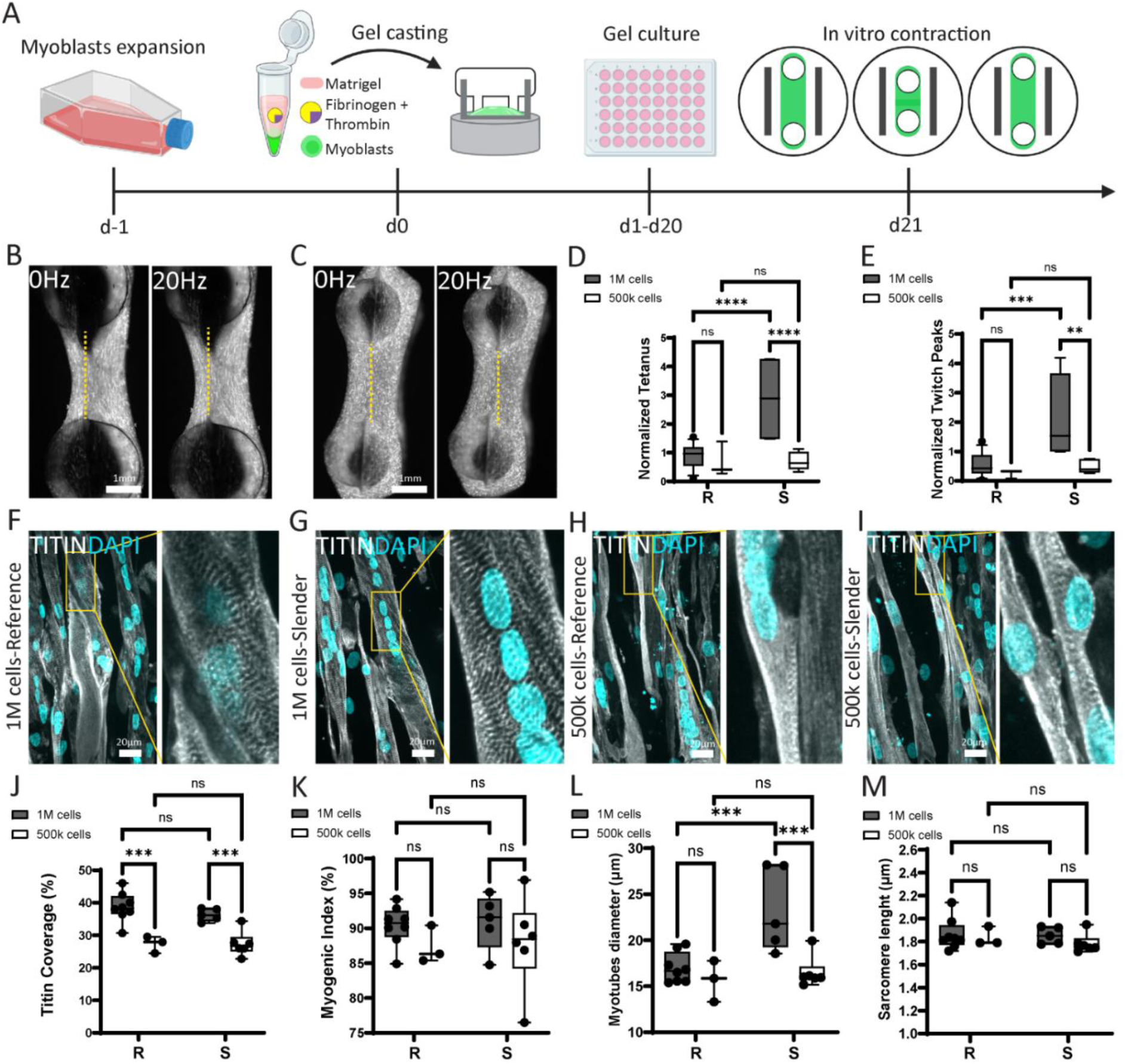
Support properties modulate function while preserving tissue maturation. (A) Schematic representation of 3D SkM generation and contraction analysis. (B, C) Representative images of tetanic force on 21 days 3D skeletal muscles generated using 1×10⁶ of SkM cells and Refence (R) or Slender (S) PDMS supports and normalized by samples area. (D, E) Quantification of tetanic and twitch forces on 21 days 3D skeletal muscles generated using 1×10⁶ or 5×10^5^ of SkM cells and R or S PDMS supports. (F-I) Representative images of 21 days 3D skeletal muscles generated using 1×10⁶ or 5×10^5^ of SkM cells and R or S PDMS supports. (J-M) Quantification of Titin coverage, myogenic index, myotubes’ diameter and sarcomere’ length of 21 days 3D skeletal muscles generated using 1×10⁶ or 5×10^5^ of SkM cells and R or S PDMS supports. Scale bars 1mm or 20μm. GFP in grey, TITIN in grey, DAPI in cyan.

### Engineered muscle tissues derived from LMNA-mutant human iPSCs cultured on PDMS scaffolds recapitulate characteristic laminopathy-associated nuclear abnormalities

Mutations in *LMNA*, encoding the nuclear lamina proteins lamin A and C, cause a spectrum of striated muscle disorders including muscular dystrophies, collectively known as muscle laminopathies. The disruption of nuclear structure and mechanotransduction resulting from these mutations leads to characteristic nuclear shape abnormalities that are among the most consistent cellular features of the disease. We and others have previously demonstrated that these phenotypes can be reproduced in hiPSC-derived engineered skeletal muscle tissues using commercially available culture platforms^23,24^. However, such systems provide limited opportunities for customization and scalable experimental design. To validate our microfabrication strategy and establish a more tuneable platform for laminopathy modelling, we investigated whether engineered muscle tissues generated on microfabricated PDMS scaffolds could recapitulate the characteristic nuclear abnormalities observed in *LMNA*-mutant muscle. We therefore evaluated whether CUSTOM (Customisable, Shape-Tuneable, Oriented Modular PDMS Scaffolds for Disease-Relevant 3D Skeletal Muscle Engineering) could reproduce key pathological features of laminopathies and serve as a reliable platform for disease modelling and therapeutic screening. Here, we combined the use of our *Reference* PDMS supports, characterized by higher stiffness, with a well-established hiPSC-derived laminopathy cell line, carrying the severe R249W pathogenic variant^25^, to recapitulate *in vitro* the characteristic nuclear morphological abnormalities associated with the disease. This approach serves as a proof of principle for future disease-focused studies using support-guided 3D skeletal muscle models. During skeletal muscle differentiation, nuclear morphology undergoes marked changes as mononucleated myoblasts fuse to form multinucleated myotubes, share a common cytoplasm, and experience increasing tensile forces associated with fiber elongation^26^. We first monitored nuclear morphological changes in 2D cell monolayers and confirmed that, in a healthy microenvironment, these nuclear changes arise as a consequence of physiological muscle maturation (Supplementary Figure 4A–H). We next investigated whether our PDMS supports could sustain and reveal nuclear morphological remodelling in 3D cultures. As described in previous experiments, we compared *Reference* and *Slender* supports using two initial cell densities (1×10⁶ and 5×10⁵ cells per hydrogel). Nuclei exhibited increased elongation in constructs cultured on *Reference* supports at the higher cell density, whereas no significant differences were detected between supports at the lower cell density (5×10⁵ cells; Supplementary Figure 4I–N). These results indicate that, in healthy cells, the stiffer *Reference* supports generate higher tension and that a cell-density threshold is required to resolve nuclear shape differences. Based on the role of mechanical tension in promoting nuclear elongation, we therefore selected the stiffer PDMS support for experiments using a laminopathy hiPSC-derived cell line. Notably, under these conditions, even the lower cell density (5×10⁵ cells) enabled the detection of disease-associated nuclear morphological alterations, when comparing control and laminopathy-derived constructs (Figure 3A, D, G, J). We then assessed maturation at two time points: days 4 and 7 of culture. Day 4 was selected as an early differentiation stage, concomitant with the onset of TITIN expression, to confirm skeletal muscle cell maturation. At the early time point, comparison between the control and laminopathy-mutated lines revealed clear differences in nuclear morphology. Control nuclei stained for Lamin A/C appeared predominantly round (Figure 3C), whereas nuclei from the mutated line were more elongated (Figure 3E) and exhibited a higher frequency of morphological abnormalities (Fig. 3F). Quantification of nuclear aspect ratio, defined as the ratio between the major and minor nuclear axes, showed a trend toward increased elongation in the mutated line; however, this difference did not reach statistical significance. When analysing the same parameters at day 7, significant differences were observed across all nuclear descriptors. Nuclei from the laminopathy mutated line displayed increased elongation, stretching, and a higher frequency of morphological abnormalities compared to controls (Figure 3H, I, K, L). These findings further support the ability of this platform to recapitulate complex skeletal muscle disease phenotypes *in vitro*.

**Figure 3.**
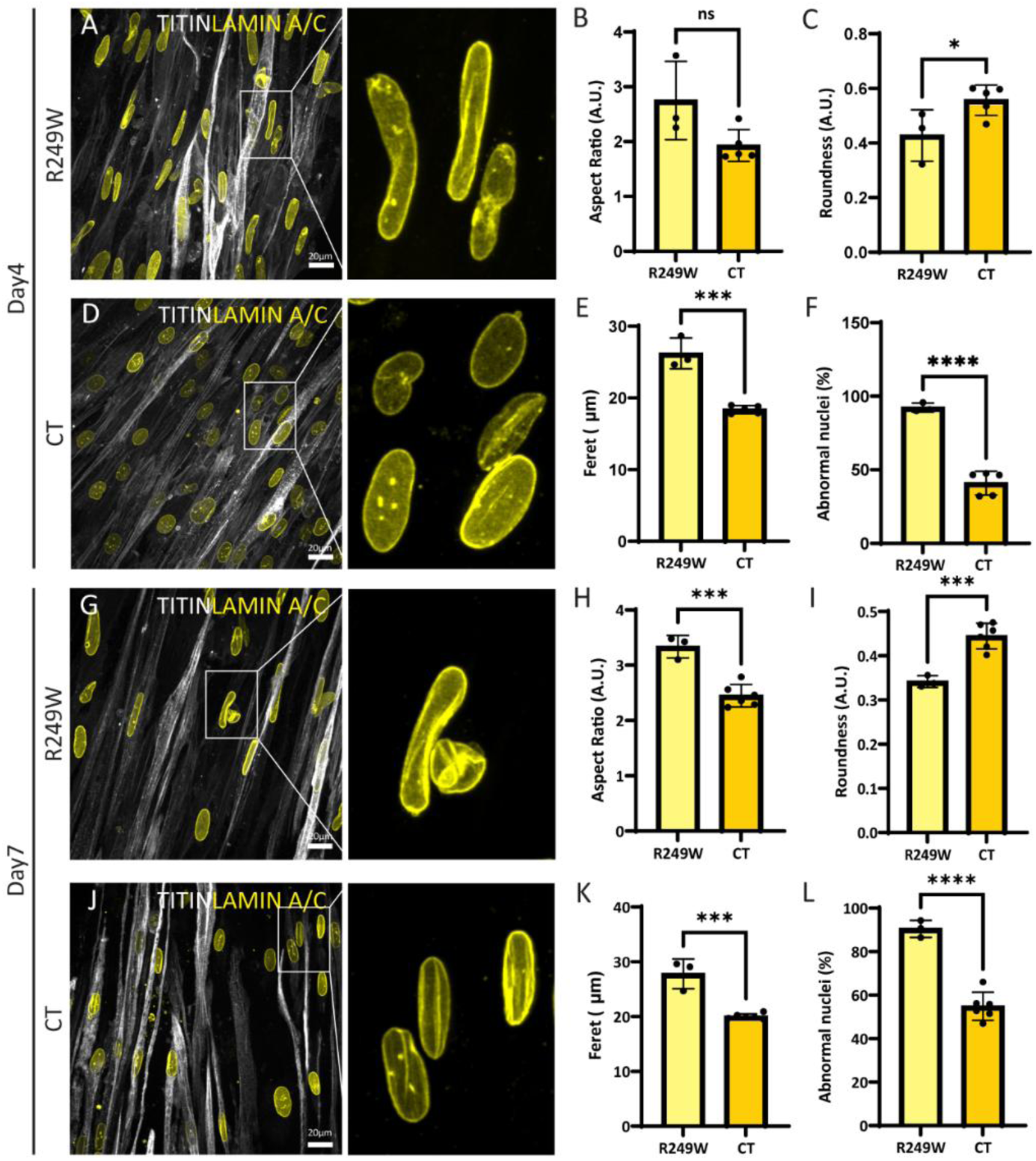
Support-guided 3D SkM hydrogels mimic the elongated nuclear phenotype of Laminopathies. (A, D) Representative images of laminopathic and control 3D SkM gels and respective quantification of nuclei elongation (B, C, E) and abnormalities (F) at day 4. (G, J) Representative images of laminopathic and control 3D SkM gels and respective quantification of nuclei elongation (H, I, K), and abnormalities (L) at day 7. Scale bars 20μm. TITIN in grey, Lamin A/C in yellow.

### PDMS-based supports facilitate motor neuron–muscle co-culture and the formation of functional human neuromuscular units

Skeletal muscle functionality strongly depends on the presence of other cell populations, which contribute not only to tissue maturation and maintenance but also to regeneration. Among these, motor neurones (MNs) play a crucial role by enabling the formation of neuromuscular junctions (NMJs), highly organized synapses between motor neurons and muscle fibres. Most *in vitro* neuromuscular models rely on incompletely matured skeletal muscle or compartmentalized culture platforms that limit the formation and functional interrogation of integrated neuromuscular tissues. Here, we adopt a modular strategy to generate a self-contained, contractile neuromuscular unit, enabling simultaneous assessment of muscle function, motor neuron–muscle interactions, and live calcium dynamics within a single platform (Figure 4A). Coupling CUSTOM muscle to MNs 3D cultures developed in our lab^9,27^ we can generate functional neuromuscular units, capable of contractility and with a complex 3D structure, which can be used not only for structural and molecular analyses post-fixation but also for real-time live phenotypic functional monitoring during culture without disturbing the samples. To achieve this, we designed dedicated adaptors that reduce and maintain a constant distance between our 3D samples and the microscope objectives, thereby enabling reliable live imaging and metabolic experiments (Supplementary Figure 5D, E). MNs were self-assembled as 3D aggregates (neurospheres, “NSs”) and evaluated for their ability to extend axonal projections within the engineered muscle microenvironment. MN aggregates were embedded within fibrin hydrogels and monitored over time for axonal outgrowth, demonstrating their capacity to adapt to the complex extracellular matrix environment of the 3D constructs (Supplementary Figure 5A). To establish neuromuscular co-cultures, MNs aggregates were subsequently seeded onto 3D skeletal muscle constructs. Following positioning on the muscle tissue, MN aggregates displayed morphological remodelling, suggesting their ability to interact with and respond to the underlying matrix and skeletal muscle architecture (Figure 4B). Taking advantage of the versatility of our system, we monitored live skeletal muscle tissues and observed continuous movement of nuclei and muscle fibers, while also confirming the formation of elongated muscle fibers (Supplementary Figure 5F). In parallel, we investigated neuromuscular interactions and confirmed not only the extension of motoneuron neurites, but also their dynamic remodelling over time, providing a real-time readout of fine contact-point formation (Figure 4C). Metabolic activity represents a crucial hallmark of cellular behaviour, and the assessment of calcium transients is particularly relevant not only in skeletal muscle alone, but especially in neuromuscular constructs. In this setup, KCl stimulation induced the expected Cav1.1/RyR1-mediated calcium influx and concurrent tissue displacement, confirming direct functional depolarisation-induced muscle contraction in the 3D tissues (Figure 4E)^28^. Furthermore, in muscle constructs with seeded neurospheres, NMDA administration caused simultaneous muscle contraction and calcium influx (Figure 4D), which was not seen in muscle constructs lacking neurospheres (Supplementary Figure 5B). Since NMDA selectively stimulates neuronal firing without directly depolarizing skeletal muscle fibers^6^, the robust calcium spikes observed exclusively in co-culture conditions confirmed the formation of functional neuromuscular junctions in our setup (Figure 4F). The maintenance of a standardized myonuclei number suggested that the neurosphere seeding approach did not negatively affect skeletal muscle tissue integrity (Figure 4I). In addition, post-fixation analysis confirmed productive engagement between the MNs and SkM fibres (Supplementary Figure 5C), with increased expression of BTX-positive spots corresponding to putative neuromuscular junctions in the areas of overlap between the two cell types (Figure 4H).

**Figure 4.**
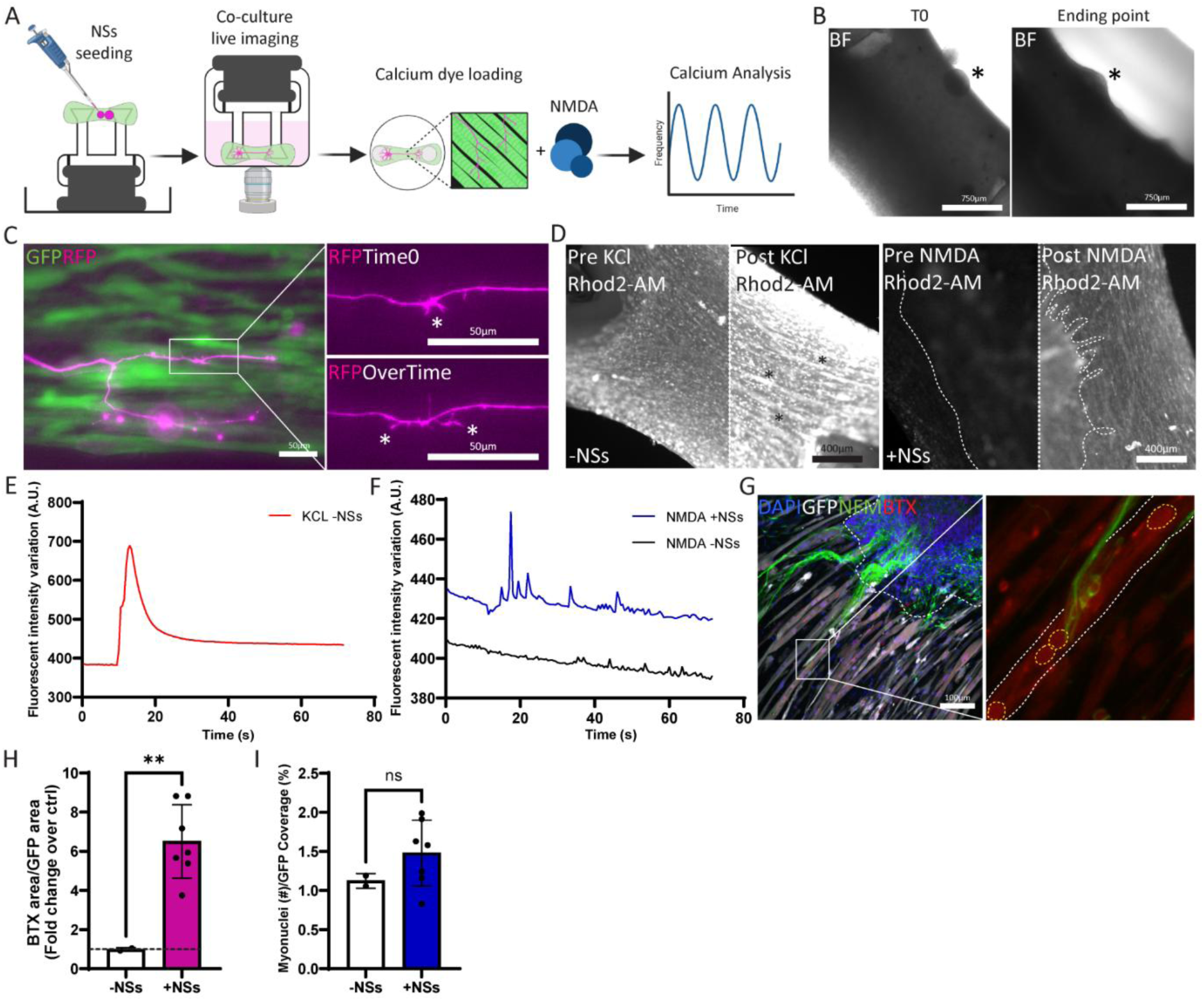
PDMS-based supports enable formation of neuromuscular constructs for high-resolution live imaging and NMJ analysis. (A) Schematic representation of 3D Neuro-SkM generation and calcium transients’ analysis. (B) Representative images of seeded neurospheres (NSs) at day 0 and 10d. (C) Representative image of neuromuscular interaction in live imaging. Magnification at time 0 and overtime of neurites sprouting. (D) Representative images of 3D neuro-SkM gels pre and post KCl or NMDA administration. (E, F) Quantification of calcium transients in 3D SkM and neuro-SkM gels post KCL or NMDA administration. (G) Representative image of 3D neuro-SkM gel and magnification of a putative NMJ spot. (H, I) Fold change of BTX spots normalized per SkM area, and quantification of the number of myonuclei per SkM with and without NSs. Scale bars 750, 50 and 100μm. Brightfield in grey, skeletal muscle in green GFP, neurites in magenta RFP, calcium in grey, GFP in grey, NFM in green, BTX in red, DAPI in blue.

### Magnetically actuated bioreactor enables *in vitro* muscle stretching, fibre disruption and tissue property characterisation

Skeletal muscle myofibres are embedded within a dynamic microenvironment that is essential for adapting to daily mechanical stress and load. This environment is often altered in skeletal muscle disorders, thereby compromising the integrity and functionality of the entire tissue. Unlike conventional scaffold-based models, which often lack customisability and integration with suitable functional analysis tools, our approach establishes an open-source, integrated ecosystem in which both devices and analytical workflows can be tailored to specific biological questions. Historically, the ability to damage skeletal muscle and study its regeneration has relied on chemically inducing injury in mature fibers. In this context, we demonstrate that a healthy 3D SkM construct (Supplementary Figure 6A, B) can be treated with cardiotoxin to induce myofiber disruption, and that this damage can be restored following myoblast injection, also within our custom-made supports (Supplementary Figure 6C–E). Beyond chemically induced skeletal muscle myofiber disruption, alternative approaches exist to investigate myofiber load and damage, including the ability to reproduce mechanical stretching *in vitro*. Based on this concept, we further explored mechanically induced stress as a complementary strategy to model muscle injury and adaptation. We designed and fabricated a magnetically actuated bioreactor capable of stretching our custom-made PDMS supports. The platform consists of a 3D-printed base housing both an electromagnet and a high-resolution camera (Figure 5A, B, Supplementary Figure 7A). A magnet is embedded within one pillar of the PDMS support (Figure 5C); when the electromagnet is activated, displacement of the pillar is induced (Figure 5E). The samples were cultured in 3D-printed, glass-sealed chambers mounted onto a custom 3D-printed holder (Supplementary Figure 7B, C). The holder and chamber assembly were connected to a commercially available adjustable stage, allowing precise regulation of the distance between the sample and the electromagnet. This setup enabled modulation of pillar displacement and, consequently, control over the degree of tissue stretching. The system was operated through a single-button interface (Supplementary Figure 7D). Stimulation frequency was controlled using a Python script; here, we present a script programmed to activate the electromagnet three consecutive times, with alternating 6-second OFF and 6-second ON periods in each stimulation cycle (Figure 5D) resulting in the intermittent displacement of the pillars (Figure 5E). Following a total of 40 stimulation cycles, the quality of the 3D SkM tissues was evaluated. Full images of the control muscles (no magnet) and unstimulated samples (magnet included but no stimulation applied) showed intact myofibers spanning from one pillar to the other. In contrast, stimulated samples (magnet included and subjected to stimulation) exhibited a clear rupture of the myofibers after 40 continuous cycles (Figure 5F). Furthermore, higher-magnification images of TITIN staining and the corresponding analysis of fiber integrity revealed marked disruption of the muscle fibers in stimulated tissues compared with controls in a cycle-dependent manner (Figure 5G, H: Supplementary Figure 8D). These findings confirm that the stimulation protocol induced substantial structural damage to the engineered skeletal muscle tissues. In parallel, the system can be used as a tool to infer muscle stiffness, opening new opportunities for disease evaluation. Taking advantage of the high degree of flexibility in support fabrication, we generated a hybrid PDMS support combining the stiffer pillar geometry of the *Reference* support with the more compliant pillar geometry of the *Slender* support. By embedding a magnet within the S pillar foot and actuating the electromagnet at different distances, skeletal muscle displacement can be measured in response to varying magnetic forces (Supplementary Figure 8A–C). These measurements provide a basis for relating muscle displacement to an externally applied force, enabling the characterization of muscle mechanical properties. This approach could therefore be used to assess and compare 3D skeletal muscle constructs with different myogenic backgrounds, providing a new tool to investigate disease-associated alterations in muscle stiffness and responses to diverse mechanical stimulation protocols.

**Figure 5.**
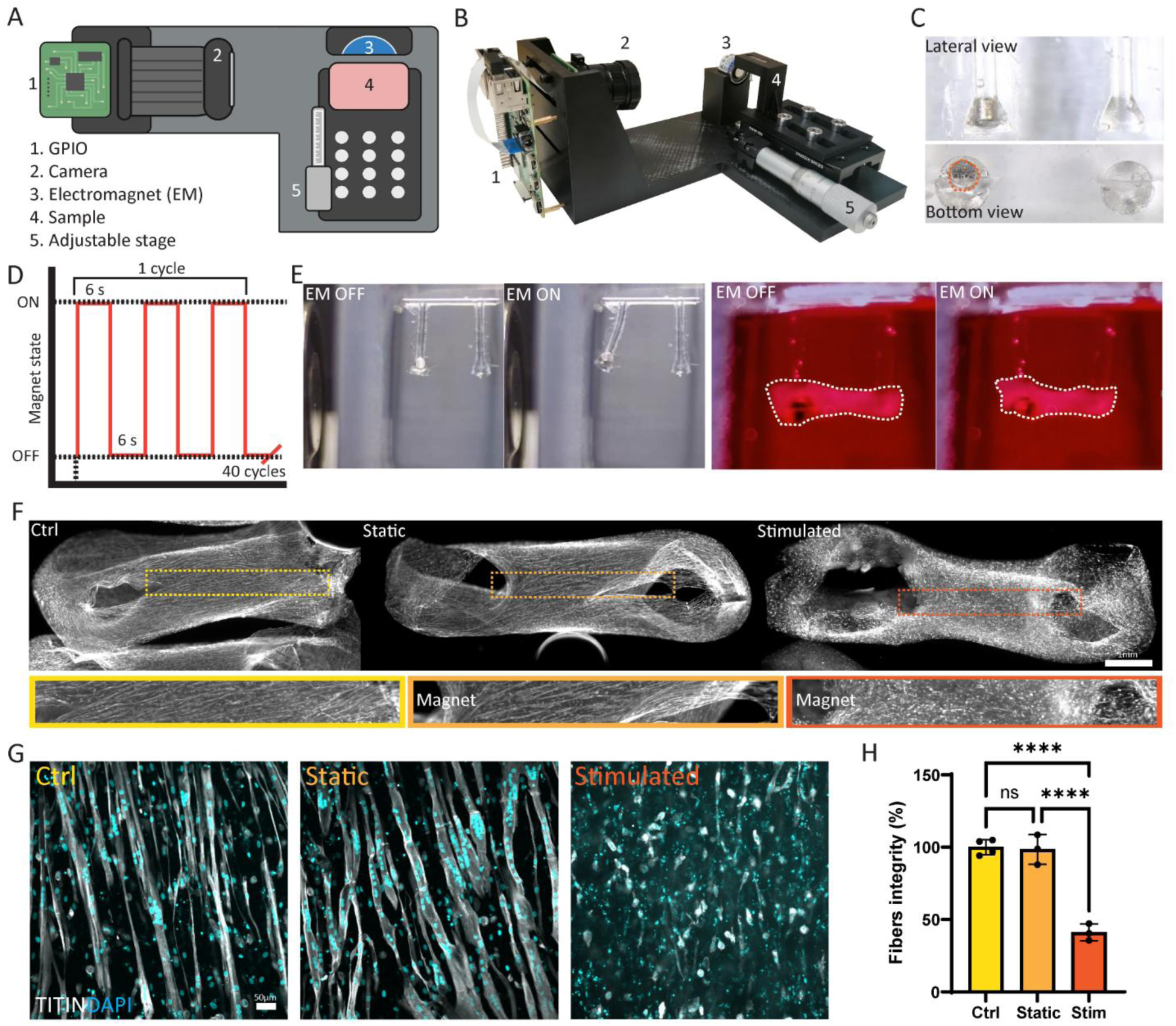
Magnetically actuated bioreactor allows long term stimulation of 3D skeletal muscle and myofibers rupture. (A, B) Schematic and representative photo of the platform. (C) Representative images of magnet inclusion in a Slender PDMS pillar. (D) Schematic of stimulation used. (E) Example of Slender pillars stimulation without skeletal muscle sample and with a 3D skeletal muscle sample. (F) Representative images of controls, magnetic unstimulated and magnetic stimulated 3D skeletal muscle gels with TITIN expression and relative magnifications on fibers’ distribution. (G, H) Representative images of control, magnetic unstimulated and magnetic stimulated 3D skeletal muscle samples and respective quantification of fibers’ integrity. Scale bars 1mm and 50μm. TITIN in grey, DAPI in cyan.

## Discussion

Gold-standard methods to engineer skeletal muscle tissue *in vitro* have predominantly relied on commercially available devices with fixed dimensions and mechanical properties, limiting their adaptability to different experimental requirements. These systems can also require relatively large cell numbers and reagent volumes, while offering limited customisation^29^. On the other hand, simplified systems, such as microfluidic devices, have significantly advanced the field by reducing cell requirements and enabling highly controlled studies^30,31^. However, these systems may offer limited opportunities to investigate tissue-level properties and integrated functional responses. Furthermore, moving to more three-dimensional organization, many existing approaches rely on fixed micropillar architectures, which often require the formation of a thin tissue layer that subsequently attaches to and wraps around two micropillars^32,33^. Such configurations can constrain tissue architecture, limit the control of muscle fibre orientation, and reduce the flexibility of tissue manipulation and experimental design. Our approach addresses these limitations by enabling each component to be customised to specific biological questions and experimental requirements, while ensuring compatibility across the platform. This modular, open-source ecosystem is designed to integrate complementary capabilities, including co-culture, live imaging, functional assessment and mechanical stimulation, paving the way towards versatile and fully integrated platforms for functional neuromuscular modelling.

Here, we presented a customisable, open-source microfabrication pipeline for generating miniaturised, physiologically-relevant, multifunctional and tuneable skeletal muscle constructs that are not restricted to a single geometry or mechanical environment, but rather can be adapted to any biological experimental pipeline, and adopted across a wide section of the field. The platform enables the rapid fabrication of customisable supports with tuneable dimensions and mechanical properties, providing a versatile framework for studying skeletal muscle development, function and disease *in vitro*.

We demonstrated that support geometry directly influences its mechanical properties, including pillars displacement, enabling the fabrication of supports with distinct mechanical environments for ad hoc experimental applications. This provides a straightforward approach to tailor the mechanical tension experienced by the developing muscle, which is essential for achieving appropriate cell alignment and maturation into functional myofibers. Importantly, mechanical properties can be adjusted to accommodate different cell lines or experimental requirements without compromising myogenic differentiation, enabling experimental conditions to be customised according to the desired biological outcome.

The three-dimensional architecture of skeletal muscle is an integral component of its complex microenvironment, which comprises a dynamic interplay of growth factors, diverse cell types, and extracellular matrix (ECM) components. Together, these elements play a crucial role in maintaining tissue homeostasis and are severely disrupted in a wide range of muscle disorders^2,34,35^. To accurately model skeletal muscle diseases *in vitro*, it is therefore important to recapitulate, as closely as possible, the three-dimensional microenvironment in which muscle cells develop and function. Although substantial efforts are still required to reproduce the different levels of tissue complexity, ranging from the development of tailored ECMs to the controlled delivery of specific growth factors, transitioning from conventional 2D cultures towards more physiologically relevant 3D environments represents an important step in disease modelling. Capturing the complexity of the native tissue organization will enable more precise experiments, support extended culture periods for longitudinal analyses, and allow the contribution of multiple factors to be investigated, features that are largely lost in conventional 2D culture systems. Here, we demonstrate that optimisation of the 3D support architecture enables to capture key cellular pathological features of a severe muscle disorders (LMNA-related muscular dystrophy). Our system not only supports efficient skeletal muscle differentiation within a complex 3D microenvironment, but also enables the detection of characteristic nuclear abnormalities and nuclear elongation associated with the disease phenotype. These findings highlight the potential of tuneable 3D skeletal muscle platforms to reproduce disease-relevant cellular phenotypes, adding an additional layer of complexity to previously published findings^23,24^ and opening new opportunities to investigate how different cell lines and disease-associated mutations influence muscle functionality *in vitro*, in a more physiologically relevant manner. Importantly, the flexibility of the platform also enables the transition from conventional longitudinal muscle geometries towards more organ-specific architectures, providing opportunities to investigate how disease-associated mutations, such as those underlying laminopathies, may differentially affect the structural and functional properties of muscle tissues with distinct anatomical organisation.

Furthermore, driven by the intrinsic need to better recapitulate tissue complexity, particularly through the incorporation of multiple cell populations, we combined our 3D skeletal muscle constructs with human iPSC-derived motor neurons. An increasing number of studies have explored the co-culture of neuronal and skeletal muscle populations, reflecting the critical role of their reciprocal interactions in supporting neuromuscular function and in the development of disease-associated phenotypes^6,36,37^ This represents an important step towards more physiologically relevant muscle disease modelling, as the ability to generate and maintain these cell populations independently before combining them provides greater flexibility in experimental design. Our supports enable the modular assembly of these distinct cellular components while allowing live imaging and straightforward sample manipulation. This flexibility opens new opportunities to investigate disease-relevant phenotypes arising from skeletal muscle–motor neuron interactions and to study neuromuscular dysfunction in a highly adaptable manner, including through the use of isogenic and allogeneic combinations of disease-associated mutations, within a more physiologically relevant 3D environment.

Finally, an important aspect of skeletal muscle tissue complexity is its ability to undergo dynamic deformation. Incorporating controlled tissue displacement *in vitro* therefore provides an additional layer of physiological relevance, enabling the mechanical microenvironment of skeletal muscle to be modulated over time. In vivo, skeletal muscle continuously undergoes cycles of damage and regeneration as part of normal tissue homeostasis. Disease-associated mutations can disrupt this balance, leading to impaired force generation, altered protein production, and progressive tissue dysfunction. However, reproducing these mechanical stresses *in vitro*, particularly in a controlled and tuneable manner, remains challenging.

We have demonstrated how the combination of our customisable PDMS supports with a newly developed 3D-printed, magnetically actuated platform enables controlled displacement of 3D skeletal muscle constructs within their native-like support environment. This system provides a flexible approach to reproducing and modulating mechanical loading *in vitro*, with the potential to be combined with mutation-specific cell lines to investigate how diseased muscle responds to mechanical stimulation. Importantly, the platform could be used not only to induce controlled tissue damage, but also to provide repeated mechanical training and to investigate the effects of prolonged mechanical loading. Such approaches could help address unresolved questions regarding the response of diseased skeletal muscle to mechanical stress, including in laminopathies and muscular dystrophies.

Despite these advances, some limitations remain to be addressed. The current manufacturing workflow relies on producing single moulds at a time, limiting scalability and throughput for larger experimental studies. Further validation using primary human skeletal muscle cells and cells derived from different muscle groups will be important to assess the platform’s versatility and its suitability for modelling muscle-specific properties. Although the current models do not fully recapitulate the native skeletal muscle niche, as they lack supporting cell populations such as satellite cells and fibroblasts, as well as a more physiologically relevant, humanised extracellular matrix (ECM), the modular nature of the platform offers considerable flexibility for further development. In particular, we anticipate that existing protocols for establishing multilineage skeletal muscle constructs using commercially available scaffolds^5,16^ can be readily adapted to this setup. Integrating these components could improve tissue maturation and better reproduce the cellular and biochemical interactions of native muscle. The platform would also benefit from the integration of high-throughput workflows and omics-based analyses to enable systematic characterisation of tissue phenotypes and molecular responses across multiple conditions. Finally, while the electromagnetic stimulation platform provides a basis for controlled mechanical loading, further development is required to reproduce more complex contraction patterns, particularly eccentric contractions, through the integration of the electromagnetic actuation system with external electrical stimulation electrodes. Addressing these limitations will be essential to increase throughput, biological complexity and functional relevance, ultimately advancing the platform towards a more comprehensive and physiologically representative system for neuromuscular disease modelling.

Overall, our approach provides a scalable and versatile platform that advances the efficiency of *in vitro* 3D skeletal muscle generation while integrating structural, cellular, and mechanical complexity. This lays a strong foundation for next-generation disease modelling, enabling the development of increasingly complex, customisable, and physiologically relevant experimental models within in-house platforms.

## Materials and methods

### Device microfabrication

3D moulds, holders, casting pockets and well adaptors were designed in Fusion 360, a computer-aided design software. Designs were then exported as STL files using Chitubox or BambuLab software. 3D moulds were printed using Sonic mini 4k resin 3D printer (Phrozen) with Aqua-Gray 4K resin (Phrozen) following our previously published protocol^9^. After printing, moulds were washed in isopropanol and UV exposed to finalise the curing process^9^. Holders, casting pockets and wells’ adaptors were 3D printed using Bambu X1-Carbon 3D printer in thermoplastic polyurethane (TPU) or acrylonitrile styrene acrylate (ASA) filaments. To obtain biocompatible skeletal muscle supports, Sylgard-184 silicone elastomer kit PDMS was used. Pre-polymer and curing agent were mixed in a 10:1 w/w ratio using a digital balance (Sartorius), and manually mixed for 5 mins prior to vacuum desiccation and casting. Demoulded PDMS was sterilized via UV exposure for 2 hours prior to use.

### Device geometry characterization

The mechanical response of PDMS pillars was first characterised analytically using Euler–Bernoulli beam theory, which models each pillar as a vertical cantilever deflected by a horizontal point load applied at its free tip. Under this framework, the maximum tip deflection is described by δ_max = FL³/3EI, where F is the applied force, L the pillar length, E the Young’s modulus of PDMS (≈ 1.5 MPa for a 10:1 crosslinker ratio^38^), and I the second moment of area of the pillar cross-section. This analytical treatment was used to explore how pillar geometry (circular, hollow cylindrical, square, or rectangular cross-sections) modulates deflection sensitivity, providing a design guide for selecting pillar dimensions appropriate to the expected force range.

### Skeletal muscle cell culture and maintenance

Human immortalized myoblasts (AB1167) were kindly provided by the Institute de Myologie (Paris, FR)^39^. Cells were transfected with a lentivirus with a GFP reporter to facilitate live visualisation. Cells were expanded in skeletal muscle cell growth medium (Promocell) at 37°C, 5% CO_2_. and passaged when 80-90% confluent using TrypLE express (Gibco). For 2D differentiation validation and nuclei morphology cells were seeded at a density of 5.000 or 10.000 cells/well. Once reached the confluency, growth medium was substituted with differentiation medium composed of DMEM high glucose (Sigma), supplemented with 1% Glutamax (Thermo), 1% Penicillin Streptomycin (Pen/Strep, Thermo) and 10 µg/ml insulin (Gibco). Control (UCLi007-A) and LMNA R249W-mutant human iPSCs (c.745C>T; https://hpscreg.eu/cell-line/CMDi004-A) were kindly provided by Cellular Dynamics International Inc. (CDI; https://www.fujifilmcdi.com/). Differentiation of hiPSCs into hiPSC-derived mesodermal progenitors (HIDEMS) was carried out as previously described^5,23,40^. HIDEMS were maintained in HIDEM proliferation medium composed of MegaCell DMEM high glucose (Sigma); 5% FBS (Life Technologies); 2mM Glutamax (Sigma); 1% Pen/Strep (Gibco); 1% non-essential amino acids; 0.05mM 2-Mercaptoethanol; 5ng/mL bFGF (Sigma)^16^. Cells were maintained in hypoxic culture conditions (5% O2, 5% CO2) with media replacement every two days. Cells were passaged when 80-90% confluent using TrypLE express.

### Motor neuron progenitors and neurospheres generation and maintenance

Wild type motor neurons (MN) were differentiated from KOLF 2.1J hiPSCs following a previously published protocol^41^. These cells were cultured in Matrigel (Corning) coated plates in a medium composed of 50% NeuroBasal (Gibco) and 50% advanced DMEM (Gibco), supplemented with B27 and N2 (Gibco), 1% Pen/Strep and 1% Glutamax. For expansion, motor neuron progenitors (MNPs) were grown and cycled with 20 ng/mL thermostable FGF2-G3 (Qkine). To generate neurospheres, MNPs were grown 1 day past confluency, then gently detached with 0.5 mM EDTA in PBS to avoid single-cell dissociation. The subsequent clumps of cells were transferred to un-coated Petri dishes and kept in suspension with 60 rpm rotary agitation. MNP spheres were grown to adequate size, then differentiation was achieved with FGF2-G3 withdrawal, and replacement with base medium supplemented with 0.1 µM compound E (Enzo), 10 ng/mL BDNF and 10 ng/mL GDNF. MNPs and neurospheres were kept in a standard humified incubation chamber at 37°C, with 5% CO_2_.

### Hydrogel casting

3D skeletal muscles were generated adapting ^9,16^. 5×10^5^ or 1×10^6^ immortalized myoblasts were resuspend in 100µL of heat inactivated skeletal muscle cell growth medium comprising 10% Matrigel, 3.5mg/mL fibrinogen (Sigma) and 3U/mL of thrombin (Baxter). The hydrogel mix was carefully pipette into ASA printed casting pockets, equipped with PDMS support. Hydrogels were left to polymerize 2 hours in humidified chamber at 37°C, 5% CO_2_. 200 µL of warm skeletal muscle cell growth medium were added on top of each hydrogel and left in humidified chamber at 37°C, 5% CO_2_ for extra 30 mins, before being carefully pipette to facilitate hydrogel removal. The 3D hydrogels attached to the PDMS supports were transferred to 48-multiwells containing 1mL of skeletal muscle cell growth medium with 33µg/mL aprotinin (Sigma). After 48 hours skeletal muscle cell growth medium was replaced with differentiation medium with 33µg/mL aprotinin. Medium was replaced every other day, hydrogels were kept in culture for 21 days, unless stated otherwise. For hydrogels containing HIDEMs, cell differentiation was induced by the addition of 1µM 4-OH tamoxifen into HIDEM proliferation medium, supplemented with 33µg/mL aprotinin. The next day medium was switched to HIDEM differentiation medium (DMEM high glucose; 2% Horse serum (Euroclone); 2mM Glutamax; 1% Pen/Strep) supplemented with 1µM 4-OH tamoxifen and 33µg/mL aprotinin. Media changes were conducted every two days and hydrogels were maintained in culture for 4 or 7 days.

For neuro-skeletal muscle hydrogels, generation, MN neurospheres were seeded on the bottom of the skeletal muscle hydrogels, 7 days after their casting. 5 neurospheres were taken in 3µL of their medium and added to other 3 µL of Matrigel. To stabilised the 3D culture, the samples were placed upside-down using ASA 3D printed well adaptors. Neuro-skeletal muscle 3D constructs were maintained for 15 mins in humified incubation chamber at 37°C, with 5% CO_2_ prior medium addition. The constructs were maintained in 48-multiwells with 1mL of skeletal muscle differentiation medium and 50/50 medium in a 1:1 ratio, supplemented with 33µg/mL aprotinin.

### Real-time visualisation of hydrogels shape

To monitor potential changes in hydrogel morphology, 3D skeletal muscle constructs were imaged on day 1 and every 7 days thereafter, up to 28 days, using the VISIBLE System^42^. Imaging was performed via the GFP channel with a 4X objective, and the entire muscle structure was reconstructed from tiled images in Fiji. Thresholder images were generated to quantify area, central width, and PDMS pillars’ distance. High-magnification images were used to assess cell orientation with the Directionality plugin in Fiji.

### Hydrogels contraction

Contractility was induced using a pair of pacing carbon electrodes (EHT-technology) in 12-multiwell containing differentiated skeletal muscle hydrogels mounted in ASA 3D printed well adaptors, enabling live visualisation of the samples and precise control of electrode spacing and stability. The stimulation consisted of bipolar square pulses at 1Hz, 100ms, 20V for twitch contraction and 20Hz for tetanic contraction. Videos were acquired via the GFP channel on an encased Nikon Eclipse Ti2 microscope with a 4X objective. The imaging chamber was humified and maintained at 37°C with 5% CO_2_. Analysis was conducted in ImageJ using the MuscleMotion plugin.

### Calcium transients

For calcium imaging experiments, neuro-skeletal muscle constructs and the respective skeletal muscle controls were incubated with Rhod-2 AM (1:1000; Invitrogen) in culture medium for 30 mins in a humidified chamber at 37°C with 5% CO₂. The 3D constructs were then transferred to ASA 3D printed well adaptors and placed in 24-well plates containing fresh differentiation medium. Videos were acquired every 500 ms using the VISIBLE System^42^. Baseline recordings were obtained prior to stimulation. To induce responses, 10 µM NMDA was applied to stimulate the neurospheres, and 100 mM KCl was used to stimulate the skeletal muscle compartment. NMDA was also applied to control hydrogels lacking neurospheres.

### Cardiotoxin damage

For chemical injury of 3D skeletal muscle constructs *in vitro*, 0.5 µM cardiotoxin (CTX) was added to the differentiation medium, and samples were incubated for 4.5 h in a humidified chamber at 37 °C with 5% CO₂. Untreated constructs cultured in differentiation medium without CTX were used as controls in each experiment. To assess the extent of CTX-induced damage, a subset of samples was fixed immediately after treatment, while others were returned to culture in fresh differentiation medium. In addition, selected constructs were injected with 2×10⁵ myoblasts 24 h after injury using 30G single-use syringes. All samples were subsequently maintained in culture for an additional 7 days, with differentiation medium supplemented with aprotinin replaced every other day.

### Immunostaining

2D and 3D experiments were fixed at different time points accordingly to the type of analysis. Cells monolayers were fixed in 4% paraformaldehyde (PFA, Boster) for 10 min at room temperature (RT) followed by three washes in 1X phosphate buffer saline (PBS, Gibco). 3D gels were fixed in 1% PFA overnight (o.n.) at 4°C and subsequently washed three times in 1X PBS. Before immunostaining, cells grown in 2D were permeabilised for 30 mins in 20mM Tris pH 7.5 solution with 150mM NaCl, containing 0.5% Triton-X100 (TBST) and then blocked for other 30 mins with TBST supplemented with 1% bovine serum albumin (BSA, Sigma), and 10% filtered foetal bovine serum (FBS, Gibco). Cells were then incubated for 1 hour at room temperature with primary antibodies, in permeabilization solution. Washed in 1X PBS and incubated with secondary antibodies or conjugated dyes, at room temperature, in permeabilization solution. Similarly, 3D gels were maintained for 6 hours in blocking solution and, after been removed from the PDMS post, incubated in mild agitation at 4°C with primary antibodies in permeabilization solution, washed in 1X PBS and incubated in mild agitation at 4°C with secondary antibodies (Invitrogen, 1:1000) or already conjugated dyes and 4’,6-diamidino-2-phenylindole (DAPI) in permeabilization solution. Samples were then mounted in cavity slides (Sigma) with FluorSave reagent (Merk). For live imaging 3D gels were incubated with SPY-DNA (Spirochrome) and Fast actin (Spirochrome) for 1 hour before removal of the dye and imaging.

### Image acquisition and analysis

Live imaging and 2D imaging were performed using an encased Nikon eclipse Ti2 with 4X and LWD 20X objectives, to allow prolonged live imaging, the chamber was humified and heated to 37°C with 5% CO_2_. 3D gels were imaged using a Nikon TiE2 CSU-W1 with 10X, 20X and 60X objectives. Z-stacks in at least 5 different gels’ regions were acquired per each sample. Myogenic index was calculated as the number of nuclei in TITIN positive myotubes divided by the total number of nuclei. Threshold images were generated to quantify TITIN and BTX clusters area, myotubes diameter was calculated in three different regions of three different myotubes per image, all data were generated using Fiji. Sarcomere length was calculated using PatternJ, a Fiij plugin previously described^43^. Nuclei morphology was calculated upon images thresholding, using Fiji shapes descriptors.

### Magnetically actuated bioreactor fabrication and operation

The bioreactor base, chamber holder, and culture chamber were designed using Fusion 360. Designs were exported as STL files and processed using Bambu Studio prior to fabrication. Bioreactor components were manufactured via 3D printing using acrylonitrile butadiene styrene (ABS) or ASA filaments. Electronic components, including a high-resolution Raspberry Pi Camera Module and a commercially available adjustable stage (Thorlabs), were integrated into the custom-built stimulation platform, which was further equipped with a 24 V electromagnet. System operation was controlled through a single-switch interface. Custom Python scripts were developed to enable live sample visualization, as well as automated image and video acquisition according to predefined stimulation protocols. Samples, casted on PDMS devices containing a neodymium magnet, were mounted within a biocompatible chamber and stimulated under differentiation medium conditions.

### PDMS supports deformation analysis

Force calibration was performed with the pillars in the free, unloaded condition. A permanent magnet embedded at the tip of one pillar was actuated by an external electromagnet, generating a reproducible magnetic load. Tip displacement was measured from two bright-field images, one with the pillar is undeflected and one with the electromagnet switched on, using a custom MATLAB pipeline. The pipeline worked in three steps. First, a pixel-to-micron scale factor was determined from a reference of known length in the undeflected image. Second, the two images were spatially registered by aligning user-defined landmarks (fixed features that do not move under magnetic actuation, such as substrate features or the pillar bases). Third, the tip position was localised by manual selection in both frames. The displacement magnitude and direction were then calculated and the applied force recovered from the Euler–Bernoulli relation with the known pillar geometry. In parallel the same magnetic actuation protocol was applied to 3D gels and measuring the resulting reduced pillar deflection allowed the stiffness of the biological construct to be estimated, since the force is now known from the prior calibration and the sample resistance is the only additional variable in the mechanical balance.

### Statistical analysis

Data are expressed as mean ± standard deviation. Statistical significance was determined with unpaired parametric Student’s t-test or ONE way or TWO WAY ANOVA, using GraphPad Prism 10. A p-value below 0.05 was considered statistically significant (*p<0.05; **p<0.01; ***p<0.001; ****p<0.0001).

## CRediT (Contributor Roles Taxonomy) Statement

EC: Conceptualization, methodology, validation, formal analysis, investigation, writing-original draft, visualization. DM: Methodology, investigation. SC: Methodology, investigation, formal analysis. FN: Methodology, investigation, formal analysis. TM: Methodology, investigation. SJ: Methodology, investigation. AI: Supervision, resources. FST: Supervision, visualization, resources. AS: Conceptualization, writing-original draft, supervision, project administration, visualization, funding acquisition.

## Acknowledgements

We would like to thank George Konstantinou and Simon Tupin for their invaluable help with the assembly of the electromagnet platform. We are grateful to the Myoline platform of the Sorbonne Université, Inserm, Institut de Myologie, Centre de Recherche en Myologie for providing the immortalised myoblast cell line used in this study.

## Generative AI statement

The authors declared that AI tools were used only for language and syntax review, and then only in an advisory role; no scientific content was AI-generated.

## Supplementary figures

**Supplementary Figure. 1.**
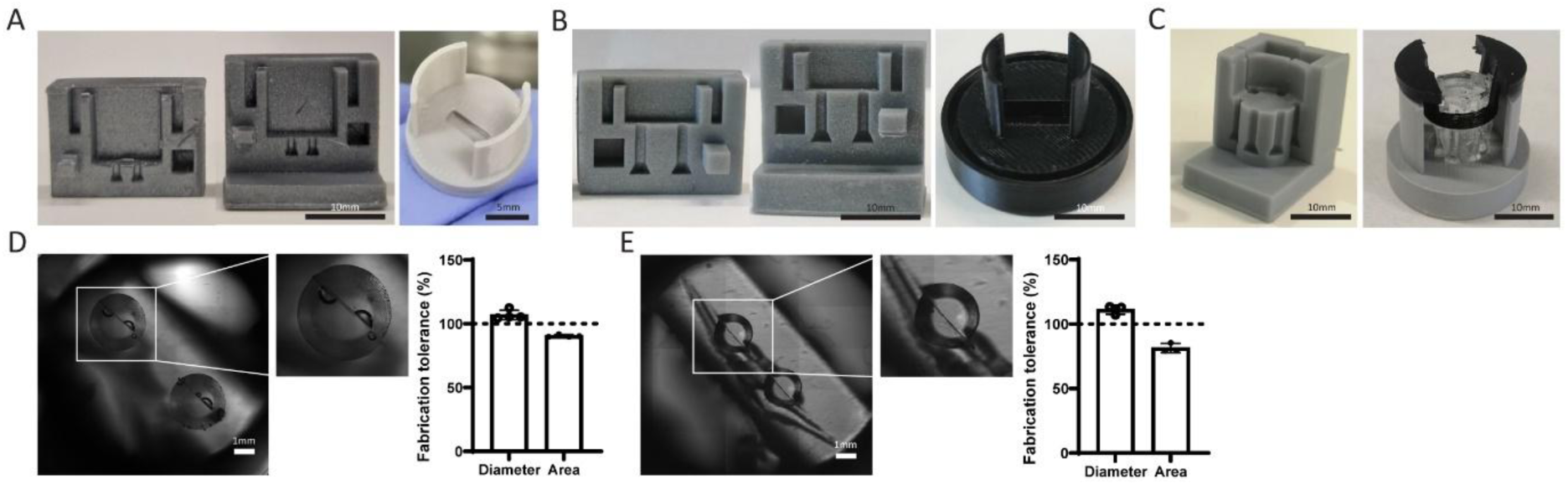
Manufacturing of 3D printed moulds and casting systems with tolerance quantification. (A, B) Resin 3D printed moulds and ASA 3D printed casting systems of M1, R and RD. (D, E) Tolerance of design and PDMS R and M1 supports. Scale bars 10, 5 and 1mm.

**Supplementary Figure 2.**
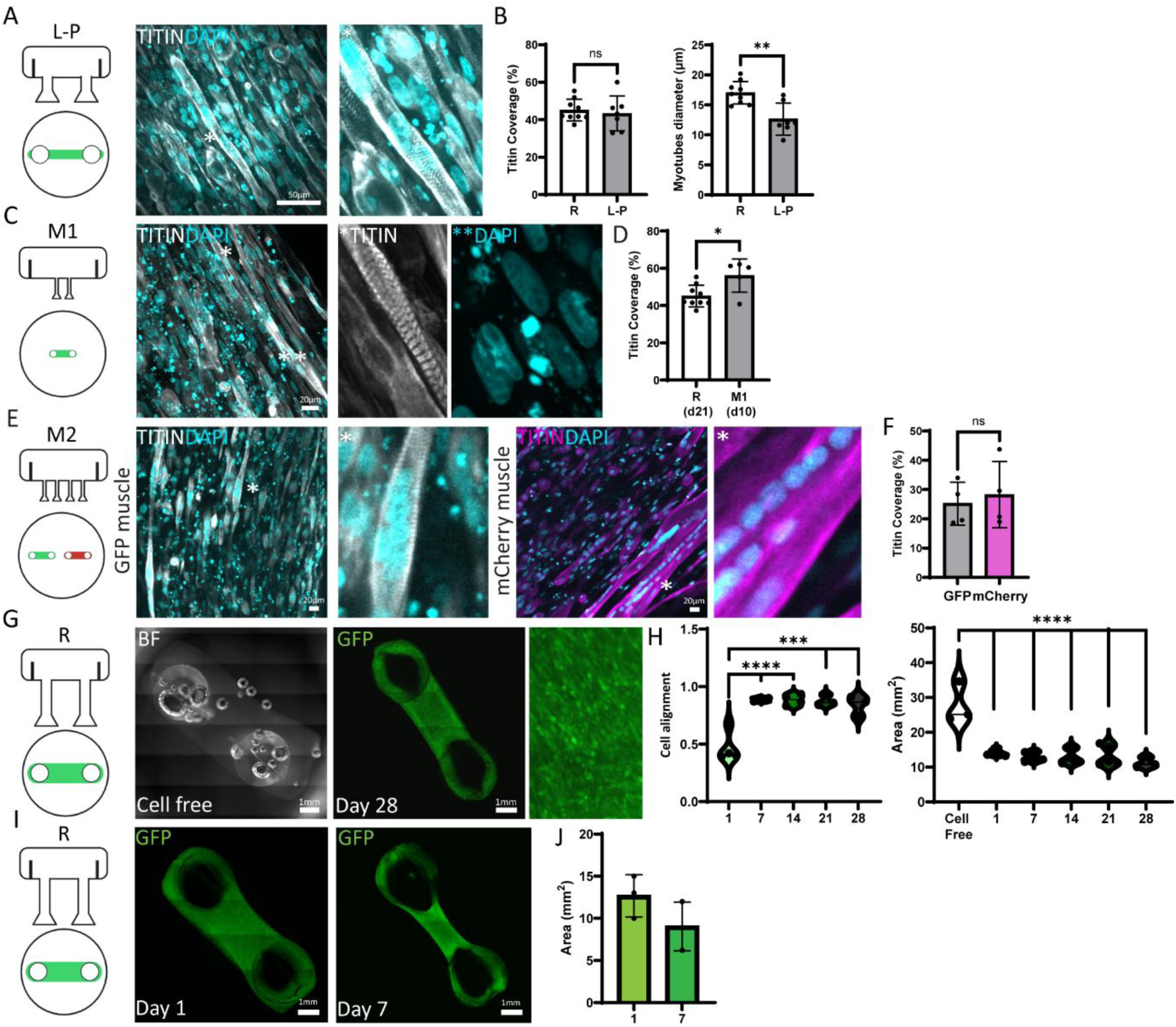
Biological validation of PDMS supports. (A) Representative image of L-P 3D SkM maturation via TITIN expression with relative magnification on muscle striation. (B) Quantification and comparison of R and L-P 3D SkM gels maturation using TITIN coverage. (C) Representative image of M1 3D SkM maturation via TITIN expression with relative magnification on muscle striation and nuclei debris. (D) Quantification and comparison of R and M1 3D SkM gels maturation using TITIN coverage. (E) Representative images of GFP and mCherry SkMs cultured on M2 PDMS support, with TITIN expression and relative magnifications. (F) Quantification and comparison of GFP and mCherry 3D SkM gels maturation using TITIN coverage. (G) Representative images of an acellular (cell free) 3D gel and of a 3D SkM gel in R PDMS supports at day 28 with respective magnification. (H) Cell alignment and gels area quantification in the 3D SkM gels at day 1, 7, 14, 21 and 28. (I) Representative images of 3D SkM gels morphology variation and respective quantification without aprotinin administration cultured in R PDMS support. Scale bars 20μm and 1mm. TITIN in grey or magenta, DAPI in cyan, GFP in grey.

**Supplementary Figure 3.**
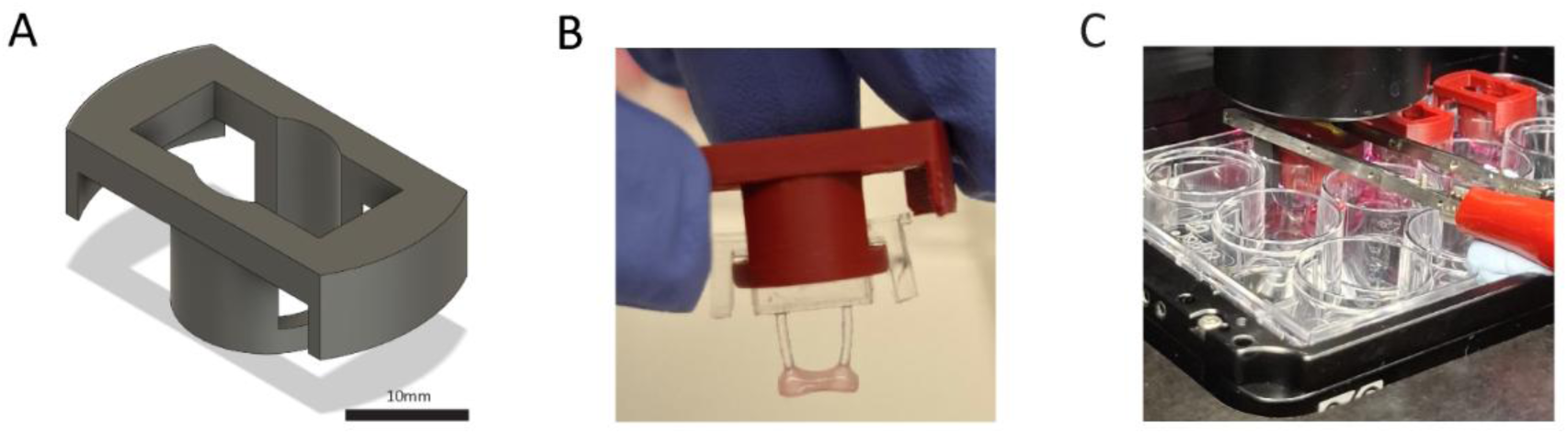
Holder design for contraction experiment. (A, B) CAD design of supports’ adaptor for contraction experiments and the respective ASA 3D printed adaptor with a mounted sample. (C) Representative photo of contraction set-up.

**Supplementary Figure 4.**
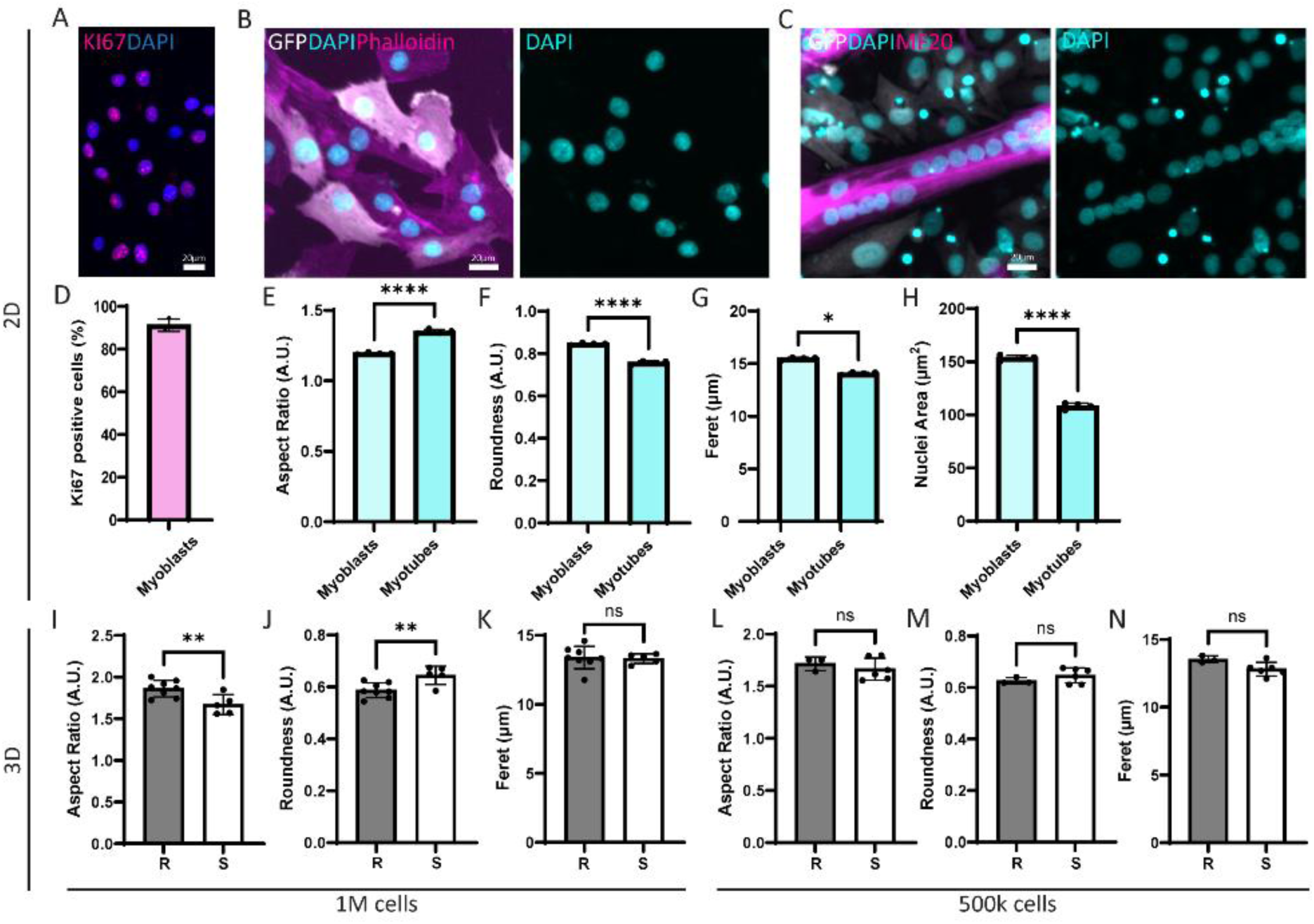
Evaluation of cell nuclei morphology in 2D and 3D microenvironment. (A, B, C) Representative images of 2D undifferentiated and differentiated myoblasts. (D) Quantification of Ki67 positive cells, (E-H) 2D myoblasts and myotubes’ nuclei morphometric analysis. (I-N) Nuclei morphometric analysis on 3D SkM gels with 1×10⁶ or 5×10^5^ cells cultured in Reference or Slender PDMS supports. Scale bars 20μm. Ki67 in magenta, GFP in grey, DAPI in cyan and Phalloidin or MF20 in magenta.

**Supplementary Figure 5.**
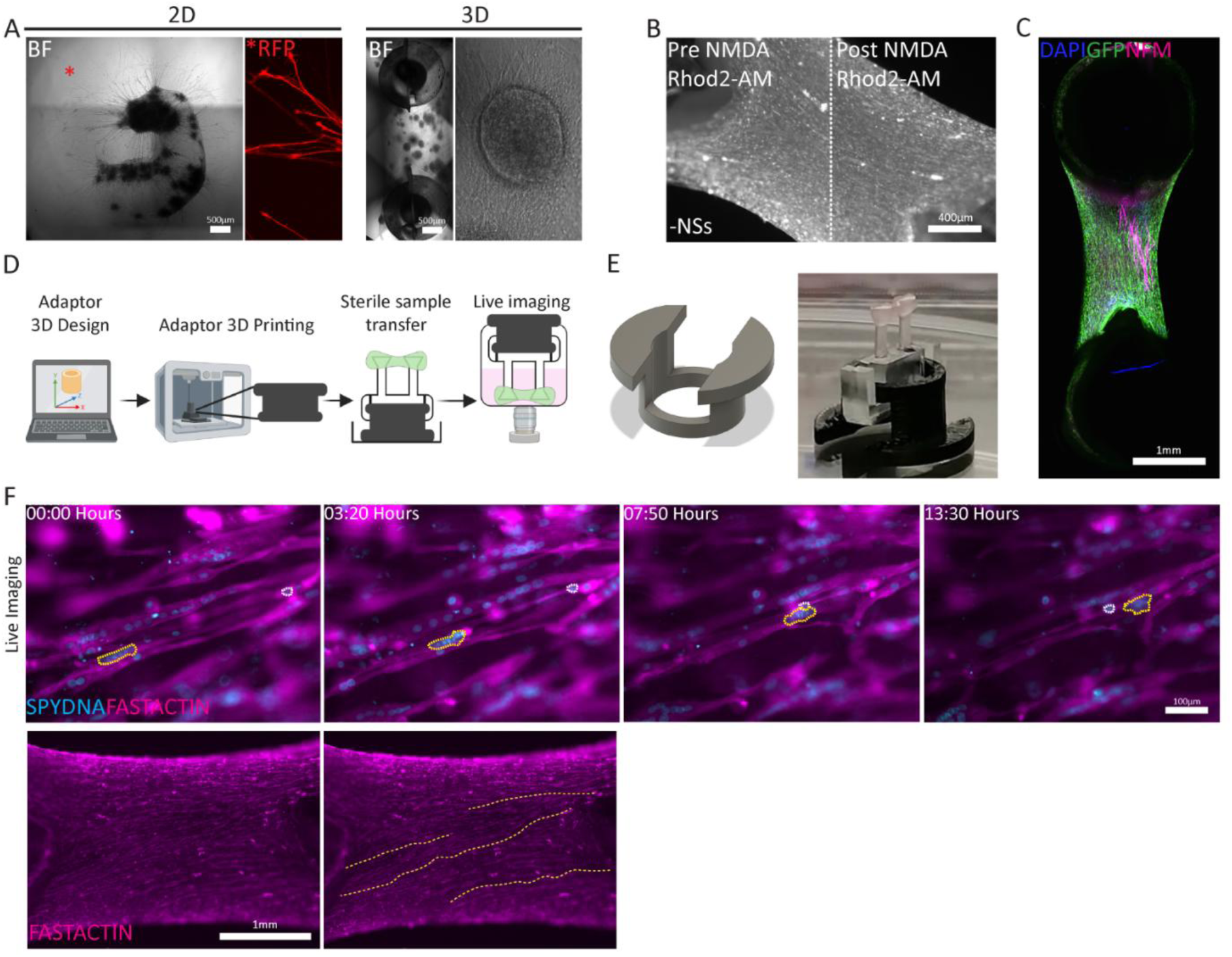
Multicellular constructs and live cell migration monitoring. (A) NSs and axonal elongation in 2D on Matrigel drops and in 3D, embedded in fibrin gel and cultured on Reference PDMS support. (B) Representative image of a 3D SkM gels before and after NMDA stimulation for calcium transients. (C) Representative reconstruction of a neuro-SkM 3D sample. (D) Scheme for the design and fabrication of a 3D SkM muscle gel adaptor converting a 48-well plate format to a 24-well plate format for precise live imaging. (E) CAD file of the adaptor and representative photo of a mounted 3D SkM muscle gel in a sterile environment. (F) Representative frames from a live imaging acquisition, of a 3D skeletal muscle generated using 5×10^5^ of SkM cells, highlighted with dotted lines the co-migration of a bag of myo-nuclei and a single nucleus and a frame showing myofibers length as possible objects of analysis. Scale bars 500, 400 and 100μm and 1mm. BF in grey, RFP in red, Rhod2-AM in grey, DAPI in blue, GFP in green, NFM in magenta, SPYDNA in cyan and Fast-Actin in magenta.

**Supplementary Figure 6.**
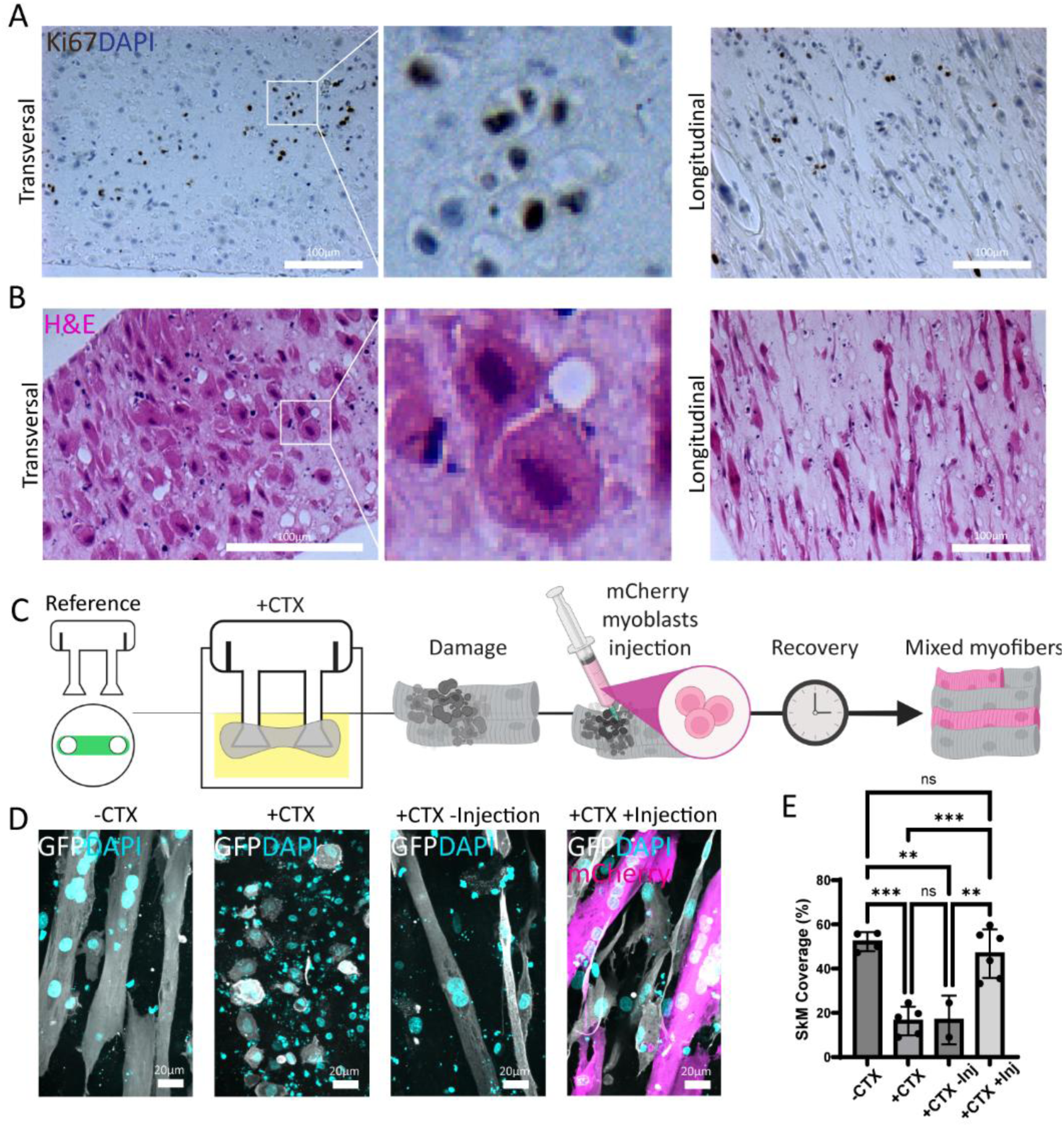
Histological validation of 3D SkM integrity and chemical CTX injury in in vitro 3D SkM. (A, B) Representative images of transversal and longitudinal cuts of 21 days 3D skeletal muscles generated using 1×10⁶ of SkM cells in Reference PDMS supports, stained with Ki67 or haematoxylin and eosin. (C) Schematic of CTX experiment. (D) Representative images of 3D skeletal muscles generated using 1×10⁶ SkM cells and reference supports, before and after CTX administration and with or without mCherry myoblasts injection. (E) Muscle cells coverage quantification. Scale bars 100 and 20μm. GFP in grey, DAPI in cyan, and mCherry in magenta.

**Supplementary Figure 7.**
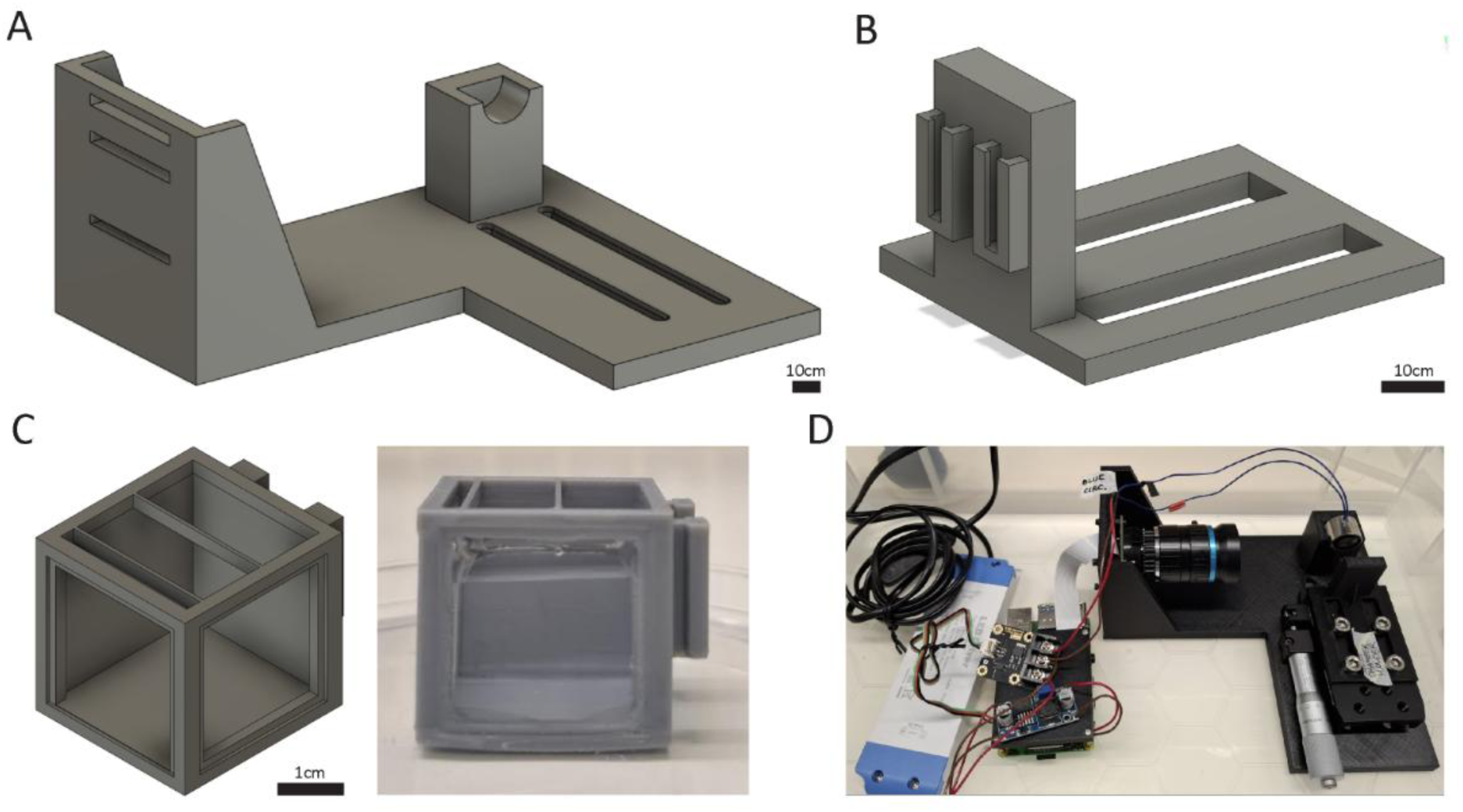
Electromagnet platform components. (A, B) CAD design of the electromagnet base and chamber holder. (C) CAD design and representative image of the sealed culture chamber ASA printed. (D) Assembled platform. Scale bars 10 and 1 cm.

**Suppl. Fig 8.**
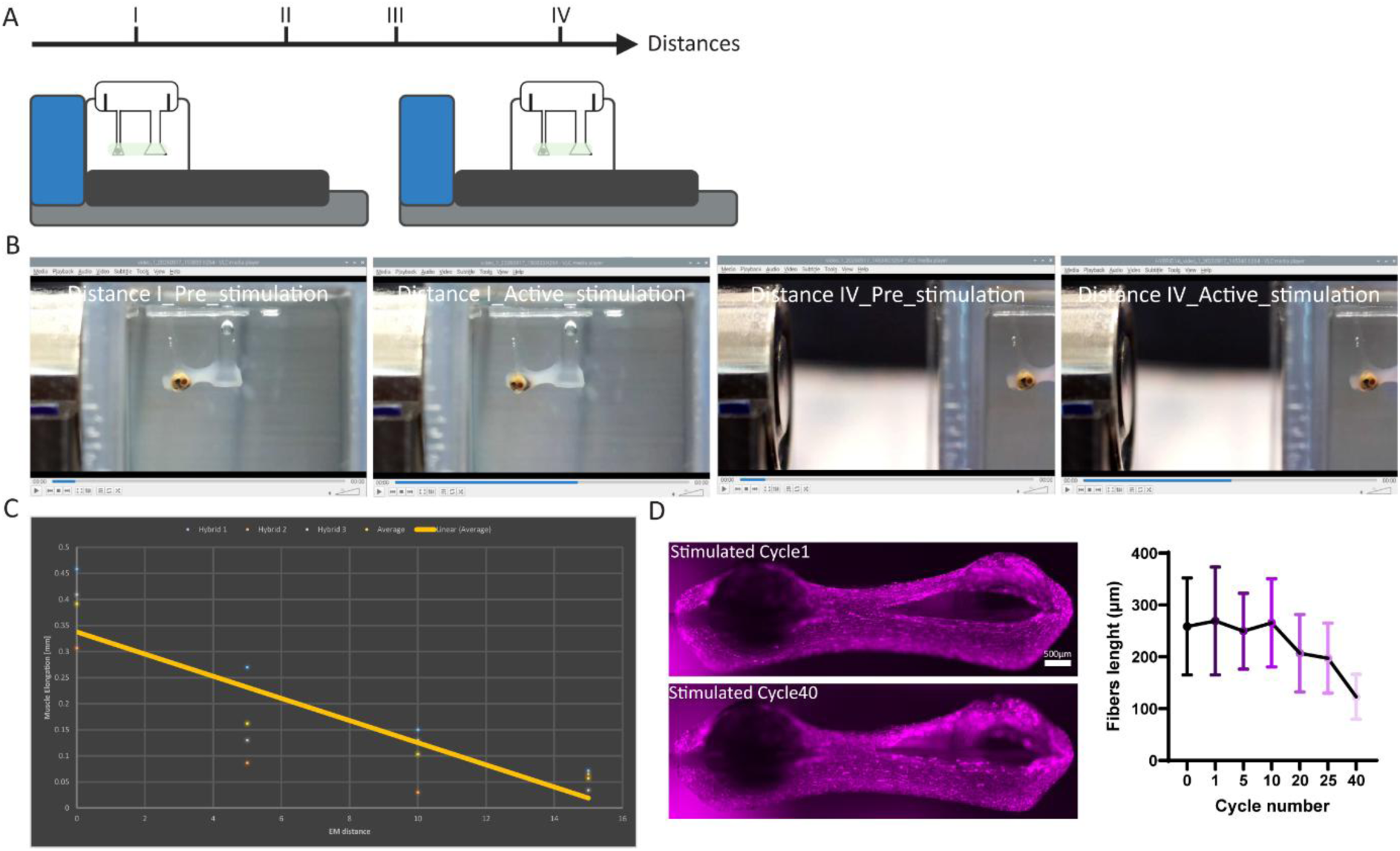
3D skeletal muscle properties and fibers tolerance. (A) Graphical scheme of the four distances used to stretch the 3D skeletal muscle cultured in hybrid PDMS supports. (B) Representative photos of 3D samples before and during active stimulation at the closest (I, 6 mm) and farthest (IV 21 mm) distances used to calculate sample stiffness. (C) Quantification of sample displacement at the four different distances, (D) Representative images of the 3D sample at cycle 1 and 40 and respective quantification of fibers rupture. Scale bar 500 μm. Fast Actin in Magenta.

## Supplementary materials

**Table 1.** PDMS devices mechanical properties.

| **PDMS Support** | **Pillar diameter (mm)** | **Pillar Radius (mm)** | **Pillar Length (mm)** | **Elastic Modulus \*(MPa)** | **Second Moment of Inertia (mm<sup>4</sup>)** | **D (mm)/F mN)** | **Ratio to Standard (R)** |
|---|---|---|---|---|---|---|---|
| **R** | 1.5 | 0.75 | 5.5 | 1.527 | 0.248504888 | 0.584591953 | 1 |
| **LP** | 1.5 | 0.75 | 2.5 | 1.527 | 0.248504888 | 0.054901573 | 0.09391435 |
| **S** | 0.75 | 0.375 | 5.5 | 1.527 | 0.015531555 | 9.353471252 | 16 |
| **Hybrid** | 0.75 | 0.375 | 5.5 | 1.527 | 0.015531555 | 9.353471252 | 16 |
| **M1** | 1 | 0.5 | 1.5 | 1.527 | 0.049087385 | 0.06003487 | 0.102695342 |

**Table 2.** Primary antibodies.

| **Antibody** | **Supplier** | **Dilution** |
|---|---|---|
| GFP | Abcam | 1:200 |
| Ki67 | Abcam | 1:100 |
| LAMIN A/C | Invitrogen | 1:250 |
| mCherry | Proteintech | 1:100 |
| MF20 | DSHB | 1:5 |
| NFM | Abcam | 1:200 |
| TITIN | DSHB | `1:11 |

**Table 3.** Conjugated dyes.

| **Dye** | **Supplier** | **Dilution** |
|---|---|---|
| BTX | Invitrogen | 1:200 |
| Fast Actin | Spirochrome | 1:100 |
| Phalloidin | Abcam | 1:1000 |
| Spy DNA | Spirochrome | 1:1000 |

